# Multiple pathways mediate chloroplast singlet oxygen stress signaling

**DOI:** 10.1101/2022.08.01.502416

**Authors:** David W. Tano, Marta A. Kozlowska, Robert A. Easter, Jesse D. Woodson

**Affiliations:** The School of Plant Sciences, University of Arizona, Tucson, AZ

**Keywords:** abiotic stress, cellular degradation, chloroplast, photosynthesis, reactive oxygen species, signaling

## Abstract

Chloroplasts can respond to stress and changes in the environment by producing reactive oxygen species (ROS). Aside from being cytotoxic, ROS also have signaling capabilities. For example, the ROS singlet oxygen (^1^O_2_) can initiate nuclear gene expression, chloroplast degradation, and cell death. To unveil the signaling mechanisms involved, researchers have used several ^1^O_2_-producing *Arabidopsis thaliana* mutants as genetic model systems, including *plastid ferrochelatase two* (*fc2*), *fluorescent in blue light* (*flu*), *chlorina 1* (*ch1*), and *accelerated cell death 2* (*acd2*). Here, we compare these ^1^O_2_-producing mutants to elucidate if they utilize one or more signaling pathways to control cell death and nuclear gene expression. Using publicly available transcriptomic data, we demonstrate *fc2, flu*, and *ch1* share a core response to ^1^O_2_ accumulation, but maintain unique responses, potentially tailored to respond to their specific stresses. Subsequently, we used a genetic approach to determine if these mutants share ^1^O_2_ signaling pathways by testing the ability of genetic suppressors of one ^1^O_2_ producing mutant to suppress signaling in a different ^1^O_2_ producing mutant. Our genetic analyses revealed at least two different chloroplast ^1^O_2_ signaling pathways control cellular degradation: one specific to the *flu* mutant and one shared by *fc2, ch1*, and *acd2* mutants, but with life-stage-specific (seedling vs. adult) features. Overall, this work reveals chloroplast stress signaling involving ^1^O_2_ is complex and may allow cells to finely tune their physiology to environmental inputs.

## Introduction

Plants experience a variety of cellular stresses, such as reactive oxygen species (ROS) produced within their energy-producing organelles (i.e., chloroplasts and mitochondria). Within chloroplasts during photosynthesis, harnessed light energy can lead to ROS production, causing damage to nearby macromolecules (e.g., lipids, proteins, DNA). Plants detoxify ROS through several enzymatic and non-enzymatic mechanisms (e.g., ROS scavenging enzymes, antioxidant production) (Apel et al. 2004; Noctor et al. 2018). However, these safety measures can be overwhelmed, especially under various environmental stresses including excess light (EL) (Triantaphylides et al. 2008), drought (Chan et al. 2016), salinity (Suo et al. 2017), and pathogen attack (Lu et al. 2018).

Photosynthesis produces the ROS hydrogen peroxide (H_2_O_2_) and superoxide (O_2_^-^) primarily at Photosystem I (PSI) and singlet oxygen (^1^O_2_) primarily at Photosystem II (PSII) (Asada 2006; Triantaphylides et al. 2008). These molecules can inhibit photosynthesis by causing photo-oxidative damage to photosynthetic machinery leading to photoinhibition (Asada 2006). Although ROS are cytotoxic molecules, they can report on a plant’s current environment (Chan et al. 2015; Foyer 2018; Mittler 2017). For instance, ^1^O_2_ can lead to signals that initiate cellular degradation (chloroplast degradation and cell death) and nuclear gene reprogramming via retrograde (chloroplast-to-nucleus) signaling (D’Alessandro et al. 2020; Dogra et al. 2019; Woodson 2022). As ^1^O_2_ has an extremely short half-life (∼0.5 – 1.0 μsec (Ogilby 2010)), the bulk of chloroplast ^1^O_2_ likely remains within the organelle of origin. Thus, secondary messengers are likely involved in propagating the chloroplast ^1^O_2_ signal(s) to affect nuclear gene expression and cellular degradation (Dogra et al. 2019; Woodson 2022). Researchers have discovered several signaling factors, but their mechanisms still remain mostly unclear.

A major challenge in deciphering ROS signaling in plant cells is that natural stresses can lead to the production of multiple types of ROS (Noctor et al. 2014; Pospíšil 2016) in multiple sub-cellular compartments (Choudhury et al. 2017; Rosenwasser et al. 2013). To specifically understand how ^1^O_2_ signals, researchers use several *Arabidopsis thaliana* mutants that conditionally accumulate chloroplast ^1^O_2_ under specific growth conditions to dissect the genetic and biochemical attributes of ^1^O_2_ signaling pathways. The *fluorescent in blue light* (*flu-1*, referred to as *flu* henceforth) mutant was one of the first ^1^O_2_-producing mutants described (Meskauskiene et al. 2001). The *flu* mutant over-accumulates the tetrapyrrole (e.g., chlorophyll and heme) intermediate protochlorophyllide (Pchlide) when grown in the dark. When the mutant is exposed to light, the energized Pchlide (like other free tetrapyrroles) reacts with nearby ground-state oxygen (^3^O_2_) to produce ^1^O_2_ within the thylakoid membranes (Wang et al. 2016). This burst of ^1^O_2_ leads to the reprogramming of hundreds of genes in the nucleus through a retrograde signal, followed by initiation of bleaching and cell death (op den Camp et al. 2003; Wagner et al. 2004). A smaller set of 168 genes, called *Early Singlet Oxygen Response Genes* (*ESORGs*), may represent the initial response (within 60 min) to chloroplast ^1^O_2_ signals in *flu* mutants (Dogra et al. 2017). A forward genetic screen for signaling components in this pathway identified *EXECUTOR1* (*EX1*) as playing an important role (Lee et al. 2007; Wagner et al. 2004). When an *ex1* loss of function mutation is introduced into the *flu* background, induction of nuclear gene expression and cell death (but not the accumulation of Pchlide or ^1^O_2_) is blocked. This breakthrough discovery was among the first evidence that ^1^O_2_-induced cell death and cellular degradation is due to a genetically encoded response rather than ^1^O_2_ toxicity.

Recent studies reveal EX1 may physically sense ^1^O_2_ in the grana margins (site of tetrapyrrole synthesis and photosystem II repair) through oxidation of tryptophan 643 in a domain of unknown function (DUF) (Dogra et al. 2019). The chloroplast metalloprotease FtsH2 degrades this oxidized EX1 protein, which is necessary for EX1 signaling (Dogra et al. 2017; Wang et al. 2016). These studies hypothesize an EX1 degradation peptide may act as a signaling factor. A conserved homolog of EX1, EX2 plays a role in ^1^O_2_ signaling (Lee et al. 2007; Page et al. 2017). Like EX1, EX2 undergoes oxidation by ^1^O_2_ at a conserved tryptophan residue and is subsequently degraded by FtsH2. However, degraded EX2 does not initiate retrograde signaling and cell death like EX1 (Dogra et al. 2022). Thus, EX2 may act as a decoy to protect EX1 and attenuate ^1^O_2_ signals to prevent excessive responses. Researchers also demonstrated, using *flu* protoplasts, the blue light photoreceptor CRYPTOCHROME 1 (CRY1) is involved in transducing the ^1^O_2_ cell death signal, leading to the possibility that blue light is involved in chloroplast stress signaling (Danon et al. 2006). However, such a signal only represents part of the ^1^O_2_ response as the impact of *cry1* on ^1^O_2_– induced nuclear gene expression is limited.

A second ^1^O_2_ over-producing mutant is *chlorina 1* (*ch1-1*, referred to as *ch1* henceforth). This mutant lacks chlorophyll b and does not have a properly functioning/assembled photosystem II antennae complex that could protect the reaction center and quench ^1^O_2_ (Ramel et al. 2013). When *ch1* is grown under EL conditions (≥ 1,100 µmol photons m^-2^ sec^-1^), PSII accumulates ^1^O_2_ in its reaction center located in the grana core. As in the *flu* mutant, the ^1^O_2_ initiates retrograde signaling to the nucleus and causes cell death. When *ch1* mutants experience a mild level of photo-oxidative stress (≥ 450 µmol photons m^-2^ sec^-1^) prior to EL treatments, they are more tolerant to subsequent EL stress, suggesting low levels of ^1^O_2_ can lead to stress acclimation (Ramel et al. 2013; Shumbe et al. 2017). In the case of the *ch1* mutant, EX1 and EX2 appear to be dispensable for ^1^O_2_ signaling (a *ch1 ex1 ex2* mutant still experiences cell death under EL stress) (Ramel et al. 2013). Instead, ^1^O_2_-triggered nuclear gene expression and cell death depends on oxidative signal inducible 1 (OXI1), a nuclear-localized serine/threonine kinase originally identified for its role in pathogen defense (Shumbe et al. 2016). Furthermore, accumulation of volatile carotenoid oxidation products (e.g., β-cyclocitral (β-cc)) produced by ^1^O_2_ accumulation at PSII are another part of this response (Ramel et al. 2012; Shumbe et al. 2014). Interestingly, signals induced by β-cc trigger nuclear gene expression and acclimation, but do not cause cellular degradation (Ramel et al. 2012). As such, we hypothesize that ^1^O_2_ signaling is a complex network controlling multiple physiological responses in the cell.

A third ^1^O_2_-producing mutant is *accelerated cell death 2* (*acd2-2*, referred to as *acd2* henceforth). This mutant experiences ^1^O_2_ bursts when grown under standard growth light conditions and produces seemingly random cell death lesions that spread across leaves (Mach et al. 2001). The *acd2* mutant accumulates the chlorophyll breakdown intermediate, red chlorophyll catabolite (RCC) (Pruzinská et al. 2007). Similarly to Pchlide accumulated in *flu* mutants, photosensitive RCC can absorb light energy and produce ^1^O_2_ in the cell (Pattanayak et al. 2012; Pruzinská et al. 2007). While the bulk (if not all) ^1^O_2_ in *flu* and *chl* mutants likely accumulates in chloroplasts (the grana margins and the grana cores, respectively), the *acd2* mutants produce at least some ^1^O_2_ in the mitochondria. Previous work did not reveal how this ^1^O_2_ may signal or lead to cell death, but this pathway acts independently of EX signaling (*acd2 ex1 ex2* mutants have similar lesion formation as *acd2*) (Pattanayak et al. 2012).

In addition to cell death and retrograde signaling, ^1^O_2_ can lead to chloroplast degradation. *plastid ferrochelatase two* (*fc2*) mutants accumulate the tetrapyrrole intermediate protoporphyrin-IX (Proto) immediately after dawn (Papenbrock et al. 2001; Woodson et al. 2015). Like Pchlide, Proto absorbs light energy and produces ^1^O_2_. The ^1^O_2_ leads to chloroplast degradation, retrograde signaling, and eventually cell death (Woodson et al. 2015). Even under permissive constant light conditions, a subset of chloroplasts (up to 35%) are selectively degraded in otherwise healthy cells, likely due to moderately high levels of Proto and ^1^O_2_ (Fisher et al. 2022). To understand the molecular signal in the *fc2* mutant, we previously conducted a forward genetic screen to identify suppressors of ^1^O_2_–triggered cell death and identified 24 *ferrochelatase two suppressor* (*fts*) mutations that allow *fc2-1* (hereafter referred to as *fc2*) seedlings to survive under diurnal cycling light conditions (Woodson et al. 2015).

When we cloned these *fts* mutants, we identified an important role for plastid gene expression in initiating the ^1^O_2_ signal in *fc2* chloroplasts. Mutations affecting *PENTATRICOPEPTIDE REPEAT CONTAINING PROTEIN 30* (*PPR30*) or “*MITOCHONDRIAL*” *TRANSCRIOPTIONAL TERMINATION FACTOR 9* (*mTERF9*) block cell death and the induction of nuclear stress genes when ^1^O_2_ accumulates in chloroplasts (Alamdari et al. 2020). These mutations lead to a broad reduction of plastid-encoded RNA-polymerase (PEP)-encoded transcripts, which is likely due to the predicted functions of PPR and mTERF proteins in post-transcriptional gene expression within plastids (Barkan et al. 2014; Wobbe 2020). In addition, we identified a third gene, *CYTIDINE TRIPHOSPHATE SYNTHASE 2* (*CTPS2*), that is necessary for ^1^O_2_ signaling in the *fc2* mutant (Alamdari et al. 2021). *ctps2* mutants are deficient in chloroplast dCTP, leading to reduced chloroplast DNA content and plastid transcripts. Based on these mutations decreasing plastid gene expression, we hypothesized that a chloroplast-encoded protein (or RNA) is essential for the *fc2* ^1^O_2_ signaling pathway (Woodson 2022).

The same genetic screen revealed the cellular ubiquitination machinery is involved with ^1^O_2_ signaling in *fc2* mutants. *FTS29* encodes the cytoplasmic plant U-box E3 ubiquitin ligase (PUB4) protein, which is necessary to induce ^1^O_2_-dependent cell death (Woodson et al. 2015). As an E3 ligase, PUB4 is likely responsible for controlling the placement of ubiquitination modifications on a group of proteins in the cell (Callis 2014). Although its targets are unknown, ^1^O_2_-stressed chloroplasts do accumulate ubiquitin-tagged proteins. Thus, PUB4 may lead (directly or indirectly) to the ubiquitination of proteins associated with the chloroplast envelope during ^1^O_2_ and photo-oxidative stress (Jeran et al. 2021; Woodson et al. 2015). Together, these conclusions suggest posttranslational modifications are a possible mechanism a cell could use to identify damaged chloroplasts for turnover or repair (Woodson 2019).

Although researchers have identified several signaling ^1^O_2_ factors, they primarily study these components in the ^1^O_2_ accumulating genetic backgrounds in which they were identified. Presently, some evidence suggests these pathways are independent; *ex1* does not suppress cell death in the *fc2* (Woodson et al. 2015), *ch1* (Ramel et al. 2013), or *acd2* mutants (Pattanayak et al. 2012). As such, we hypothesize that multiple chloroplast ^1^O_2_ signaling pathways exist to control cellular degradation and nuclear gene expression. Here, we compare the *fc2, flu, ch1*, and *acd2* mutants to test if they elicit separate chloroplasts signals. A meta-analysis of transcriptional responses in these mutants suggests a core response with unique signatures exists. However, a genetic analysis of these mutants and their suppressors revealed two major ^1^O_2_ signaling pathways. The *flu* mutant likely uses one distinct EX-dependent signal, while *fc2, ch1*, and *acd2* share a second ^1^O_2_ signaling pathway with life-stage-specific (seedling vs. adult) features. Together these results demonstrate chloroplast ^1^O_2_-signaling is complex and may depend on the exact sites of ^1^O_2_ production, even within a single chloroplast.

## Methods

### Biological material, growth conditions, and treatments

The Arabidopsis ecotype *Columbia* (Col-0) was used as wt and the genetic background for all lines. Mutant lines used in this study are listed in Table S1. The *fc2-1* T-DNA insertion line (GABI_766H08) (Woodson et al. 2011) and the *oxi1* T-DNA insertion line (GABI_355H08) (Camehl et al. 2011) used were from the GABI (Kleinboelting et al. 2012) T-DNA collections and were described previously. The *ex1* (SALK_002088), *ex2-2* (SALK_021694), and *ex2-3* (SALK_121009) T-DNA insertion lines used were from the SALK T-DNA collections (Alonso et al. 2003). SALK_002088 (Lee et al. 2007) and SALK_021694 (Uberegui et al. 2015) were previously described. The *cry1-304* (Bruggemann et al. 1996), *pub4-6* (Woodson et al. 2015), *acd2-2* (Mach et al. 2001), *flu-1* (Meskauskiene et al. 2001), *ppr30-1* (Alamdari et al. 2020), and *ch1-1* (Havaux et al. 2007) mutants were described previously. Higher order mutant combinations were generated by crossing and confirmed by PCR genotyping where applicable (primer sequences listed in Table S2).

Seeds were surface sterilized and plated using one of two methods; 1) a previously described liquid bleach washing protocol (Alamdari et al. 2020). Briefly, seeds were washed in 30% bleach with 0.04% Triton X-100 (v/) and then rinsed with sterile water three times by pelleting seeds at 3,500 x g for 1 min. 2) Chloride gas sterilization. For chloride gas surface sterilization, approximately 25-100 μl of seed was placed in 2 mL microcentrifuge tubes and placed in an airtight chamber with their lids open. Five mL of concentrated HCl were added to 150 mL of bleach (3.33% v/v) and the lid to the chamber was put on immediately. Seeds were removed 24 h later and allowed to air out for 15 min before plating. Sterilization Method 1 was used for plants monitored or assayed in the seedling stage. Sterilization Method 2 was used for growing plants for bulking seed, genotyping, and adult-stage experiments. Seeds were plated on Linsmaier and Skoog medium pH 5.7 (Caisson Laboratories North Logan, UT) with 0.6% micropropagation type-1 agar powder. Seeds were stratified for 4 to 5 days in the dark at 4°C and were germinated in control conditions: constant white light (∼120 µmol photons m^-2^ sec^-1^) at 22°C. To initiate stress signaling in *fc2* mutant seedlings, plates were germinated under 6 h light / 18 h dark diurnal light cycling conditions. To initiate cell signaling in *flu* seedlings, plates were germinated under control conditions, shifted to the dark after 5 days for up to 24 h (by wrapping in aluminum foil), and re-exposed to light.

To test adult phenotypes, seedlings were grown under seedling control conditions, transferred to soil, and grown under adult control conditions (100 µmol photons m^-2^ sec^-1^ of constant light at 22°C). To initiate stress signaling in *fc2* and *flu* adult mutants, plants were shifted to 16 h light / 8 h dark diurnal light cycling conditions when 21 days old. Seeds used for experiments were harvested from plants of a similar age. Photosynthetically active radiation was measured using a LI-250A light meter with a LI-190R-BNC-2 Quantum Sensor (LiCOR). All above experiments were performed in chambers with cool white fluorescent bulbs.

To initiate and monitor EL signaling in seedlings, plants were grown as described above, but in a Percival LED-30L1 with white LED lights at 120 µmol photons m^-2^ sec^-1^. When 7 days old, the seedlings were then transferred to an EL chamber (a Percival LED 41L1 chamber with SB4X All-White SciBrite LED tiles) and exposed to 1,200 µmol photons m^-2^ sec^-1^ white light at 10°C for 24 h. Maximum PSII quantum yield (F_v_/F_m_) was monitored in a FluorCam chamber (Closed FluorCam FC 800-C/1010-S, Photon Systems Instruments) as previously described (Lemke et al. 2021). For adult plants, seven-day-old seedlings were transferred to soil and grown under 70 µmol m^-2^ sec^-1^ white light from LED panels until plants were 18 days old. The plants were then exposed to 1,300 µmol photons m^-2^ sec^-1^ white light at 10°C in the EL chamber. F_v_/F_m_ was monitored the same as for the seedlings.

### Transcriptome Data Analysis

Previously published microarray expression data was gathered from studies describing *fc2* (Woodson et al. 2015), *flu* (op den Camp et al. 2003), *ch1* (Ramel et al. 2013), and ß-cc treated wt plants (Ramel et al. 2012). RNAseq data of *flu* mutant seedlings to identify ESORGs was from (Dogra et al. 2017). As the Affymetrix GeneChip Arabidopsis ATH1 Genome Array (*fc2* and *flu* datasets, Table S3) and the CATv5 microarray (*ch1* and ß-cc datasets, Table S4) have different gene coverage, only genes contained in both were used in the analysis. Gene groups recognized by a single probe were also removed as expression values could not be assigned to one specific gene. Finally, organellar gene transcript levels were removed from the analyses and only nuclear-encoded transcripts were considered. This left a total 19,895 genes for comparative analyses (Table S5).

Differentially expressed genes (DEGs) (induced or repressed) were identified from each dataset. For the *fc2* dataset, four-day-old etiolated wt and *fc2* seedlings were compared 2 h after light exposure (Table S6). For the *flu* dataset, wt and *flu* adult plants were compared after 8 h dark and 1 h light re-exposure (Table S7). For the *ch1* dataset, *ch1* adult plants were treated with 2 days of EL and compared to untreated *ch1* (Table S8). For the ß-cc treatment dataset, wt plants treated with ß-cc for 4 h were compared to water-treated controls (Table S9). For these datasets, we applied cutoff values of ± ≥ 2-fold mean expression and adjusted (Bonferroni) p-value ≤ 0.05. However, for the adult *flu* mutant dataset (op den Camp et al. 2003), an unreported significance cutoff was already applied by the authors. For the ESORG dataset, *flu* seedlings were placed in the dark for 4 h and then exposed to 30 or 60 min of re-illumination and compared to *flu* seedlings without re-illumination. A list of 168 ESORGs were identified (≥ 2-fold induction, FDR ≤ 0.05 cutoffs) that overlapped with an earlier analysis of induced transcripts in the *flu* mutant (Chen et al. 2015) (Table S10). These gene lists were then compared using the program Venny 2.1 by Juan Carlos Oliveros (https://bioinfogp.cnb.csic.es/tools/venny/index.html) (Oliveros (2007-2015)). Genes overlapping between mutants are listed in Tables S11 (up-regulated) and S12 (down-regulated), between mutants and ß-cc treatment are listed in Tables S13 (up-regulated) and S14 (down-regulated), and between mutants and ESORGs are listed in Table S15. Table S16 displays additional details regarding plant growth, and RNA extraction/processing to generate the published datasets.

### Gene ontology enrichment analyses

Using gene lists from Tables S11-15, gene ontology (GO) terms were identified using GO::TermFinder (https://go.princeton.edu/cgi-bin/GOTermFinder) (Boyle et al. 2004) and GO terms were selected based on a p-value ≤ 0.01. Qualifying GO terms were exported to REVIGO for visualization (http://revigo.irb.hr) (Supek et al. 2011).

### Polymerase chain reactions and genotyping

Approximately 100 mg of fresh tissue was flash-frozen in liquid nitrogen for five min and crushed using 2 silica beads in a Mini-BeadBeater (Biospec Products) for 1 min in 2 mL microcentrifuge tubes. DNA was extracted using 750 µL of 2xCTAB (2% w/v) solution (1.4 M NaCl, 100 mM Tris-Cl pH 8.0, 20 mM EDTA) with 0.3% v/v beta-mercaptoethanol. Samples were incubated for 20-24 h at 65°C. Debris was pelleted for 5 min at 10,000 x g at room temperature. 700 uL of the supernatant was moved to a clean tube and a 1:1 chloroform extraction was performed. Tubes were mixed for 2 min and left to rest for 5 min. Next, tubes were centrifuged for 10 min at 10,000 x g to separate the aqueous and organic phases. 600 µL of the aqueous layer was moved to a new tube containing 240 µL 5 M NaCl and 840 µL of 100% isopropyl alcohol. Tubes were mixed for 2 min, incubated at room temperature for 10 min, and incubated at 4°C for 24 h. DNA samples were pelleted for 30 min at 4°C at 21,000 x g. Two 1 mL 75% ethanol washes were performed, mixing the tubes by hand and pelleting for 2 min at 4°C at 21,000 x g, pouring the supernatant off each time. The tubes were spun at 21,000 x g for 30 seconds and the supernatant was removed. DNA pellets were dried for 2 h in a laminar flow hood and resuspended in 75 µL of DNAse-free water, incubating for 24 h at 4°C.

PCR samples were amplified using GoTaq Green Master Mix (Promega) according to the manufacturer’s instructions. 20 µL reactions were performed, using 10 µL of GoTaq Green Master Mix, 1 µL of 10 µM primer A, 1 µL of 10 µM primer B, 6 µL of sterile water, and 2 µL of genomic DNA sample. For PCR samples not requiring restriction enzyme digestion (see below), DNA fragments were separated in a 1% (w/w) agarose gel containing 0.625 mg/mL ethidium bromide for 30 min at 120 volts. Gels were imaged using a UV box. For unknown reasons, we were unable to amplify the left border of the *oxi1* T-DNA (*GABI_355H08*) using primers specific to the left T-DNA border and the *OXI1* sequence (JP1291/JP285). Instead, this mutation was confirmed by the inability to amplify wt *OXI1* using the primer set JP1291/JP1292 and 100% resistance (no segregation) to 5 μg/ml sulfadiazine (the antibiotic marker cassette in GABI T-DNA sequences).

For genotyping requiring a restriction enzyme digestion (dCAPs), 10 µL digestions were performed. In a new tube, 5 µL of PCR product, 4.4 µL of nuclease-free water, 0.5 µL of the appropriate 10x buffer, and 0.1 µL of the appropriate enzyme were combined and mixed gently by hand. Samples were incubated at 37°C overnight. DNA fragments were separated in a 3% agarose gel containing 0.625 mg/mL ethidium bromide until the dye front was at the end of the gel. The gel was imaged using a UV box. Table S2 lists enzymes and expected fragment sizes.

### RNA extraction and Real-time quantitative PCR

Total RNA extraction, cDNA synthesis, and RT-qPCR was performed as previously described (Alamdari et al. 2020), using the RNeasy Plant Mini Kit (Qiagen), Maxima first strand cDNA synthesis kit for RT-qPCR with DNase (Thermo Scientific), and the SYBR Green Master Mix (BioRad), respectively, according to the manufacturers’ instructions. RT-qPCR experiments were all performed using a CFX Connect Real Time PCR Detection System (Bio-Rad). For expression analyses, all genes were normalized using *ACTIN2* as a standard. The primers used for RT-qPCR are presented in Table S2.

### Chlorophyll measurements

Chlorophyll was measured as previously described (Alamdari et al. 2021). Briefly, seeds were stratified for 5 days and counted prior to germination. Seedlings were collected 7 days after germination. Approximately 30-60 seedlings were used per seed line, and at least 3 biological replicates were collected for both constant light and diurnal cycling light conditions. Seedlings were flash-frozen in liquid nitrogen for 5 min and crushed using a Mini-BeadBeater (Biospec Products) for 1 min. Constant light samples were extracted in 1.2 mL of 100% ethanol, and diurnal light samples were extracted in 0.150 mL of 100% ethanol. Cell debris was pelleted and removed at 12,000 x g for 30 min at 4°C. The debris removal process was repeated twice before readings were taken. Chlorophyll was measured spectrophotometrically at 652 nm and 665 nm with a Varioskan LUX spectrophotometer with optically clear 96 well plates. Path corrections were calculated and chlorophyll concentrations were determined based on a previously described protocol (Warren 2008). Each biological replicate is a mean of 3 technical replicates. Total chlorophyll content was normalized to the number of seedlings collected.

### Protochlorophyllide measurements

Protochlorophyllide (Pchlide) was measured as previously described (Shin et al. 2009). Briefly, seeds were stratified for 4 days and counted prior to germination, which was initiated with 1 h of white light in control conditions. Seedlings were grown in the dark for 4 days at 22°C. Tissue (10 seedlings per replicate) was collected in dim green light and stored in amber 1.5 mL tubes containing 2 silica beads after flash-freezing with liquid nitrogen. Seedlings were crushed using a Mini-BeadBeater (Biospec Products) for 1 min. Pchlide was extracted using 1 ml of 80% acetone (v/v). In a black plastic 96-well plate (Grenier Bio-One), 200 μl of sample was loaded, with 3 biological replicates per genotype. The fluorescence of the samples was measured (excitation 440 nm/emission at 638 nm) with a Varioskan LUX spectrophotometer.

### Singlet oxygen measurements

Singlet oxygen was measured as previously described (Alamdari et al. 2020). Briefly, seedlings were grown on plates in 6 h light / 18 h dark diurnal cycling light conditions. As day three concluded, seedlings were moved to 1.5 ml microcentrifuge tubes containing 250 μl of ½-strength Linsmaier liquid media, wrapped in foil, and incubated at 22°C for 18 h in the dark. An hour prior to subjective dawn on day four, 50 μM of 1.5 mM Singlet Oxygen Sensor Green (SOSG, Molecular Probes) and 0.1% Tween 20 (v/v) was added to the medium under dim, green light (final concentration of 250 μM). Seedlings were vacuum infiltrated for 30 min in the dark. After 30 additional min, seedlings were exposed to light for 3 h. Seedlings were washed once with 1 ml of ½-strength Linsmaier and Skoog medium pH 5.7 prior to imaging with a Zeiss Axiozoom 16 fluorescent stereo microscope equipped with a Hamamatsu Flash 4.0 camera and a GFP fluorescence filter. At least 12 seedlings from each genotype were monitored and average fluorescence per mm^2^ was quantified using ImageJ, choosing the brightest cotyledon per seedling.

### Assessment of cell death

Cell death was measured in plant tissue as previously described (Woodson et al. 2015). Briefly, tissue was stained with a trypan blue solution (10 ml phenol, 10 ml glycerol, 10 ml lactic acid, 10 ml H_2_O, and 0.02 mg trypan blue (Sigma)) diluted with 2 volumes of 100% ethanol. The tissue in the staining solution was boiled for 2 min at 100°C and incubated at room temperature overnight. Non-specific stain was removed using 2 overnight chloral hydrate (25 g / 10 mL water) incubations. Tissue was moved to 30% glycerol for imaging. The intensity of the trypan blue stain was measured with ImageJ using at least 6 seedlings from each genotype and was normalized to the area of the cotyledon and then wt. The darkest cotyledon per seedling was chosen for measurements.

### Lesion Counting

To assess leaf lesion formation in adult plants, plants were grown under 16 h light / 8 h dark diurnal cycling light conditions until lesions became apparent in some plants (day 18). Lesions were counted for each plant for an additional 18 days.

## Results

### Singlet oxygen accumulation in the *fc2, flu*, and *ch1* backgrounds leads to overlapping nuclear transcriptomic responses

The Arabidopsis *fc2* (Woodson et al. 2015), *flu* (op den Camp et al. 2003), and *ch1* (Ramel et al. 2013) mutants produce excess ^1^O_2_ in the chloroplast, which leads to the induction of nuclear genes and cell death. However, we do not know if these three mutants utilize the same ^1^O_2_ signaling pathways to promote these outcomes. To test if these mutants share ^1^O_2_ pathways, we assessed the similarity of the nuclear responses to chloroplast ^1^O_2_ accumulation. We mined publicly available gene expression datasets to identify targets of the ^1^O_2_ signal in each genetic background (i.e., differentially expressed genes (DEGs)). For the *fc2* dataset, four-day-old etiolated (dark-grown) seedlings were exposed to light for 2 h (Woodson et al. 2015). The *flu* dataset was generated using plants grown to the rosette life stage, incubated in the dark for 8 h, and then collected 1 h post light re-exposure (op den Camp et al. 2003). Finally, the *ch1* dataset was from plants grown for 5-8 weeks and exposed to 8 h of EL for 2 days (Ramel et al. 2013). We also analyzed datasets from β-cc-treated wt plants. These plants were grown for 4 weeks and exposed to β-cc for 4 h before sample collection (Ramel et al. 2012). Finally, we compared the list of ESORGs identified in *flu* seedlings grown for 5 days under constant light, dark incubated for 4 h, and re-exposed to light for 30 or 60 min before sample collection (Dogra et al. 2017). Unfortunately, there is not a publically available transcriptome dataset for the *acd2* mutant. Additional information on the datasets is listed in Table S16.

Next, we filtered the datasets to identify DEGs using cutoffs of ≥ 2-fold difference and a p-value ≤ 0.05 (Tables S6-10). Finally, we compared these lists of genes for each background/treatment group to identify DEGs shared between groups (Tables S11-15). A comparison of the identified 1,633 DEGs showed for each mutant, the majority of upregulated and downregulated DEGs were unique (Figs. 1A and B, Table S17). At the same time, a subset of DEGs were shared between the mutants (between 28.8-40.9% of total DEGs from one mutant overlapped with another). While the overlap appeared similar between the three mutants (8.5–31.0% DEGs within a set), the overlap between the *flu* and *ch1* DEGs was slightly larger (31.0% of *flu* DEGs and 23.1% of *ch1* DEGs) than with *fc2* (13.8% and 8.5% of *flu* and *ch1* DEGs, respectively). In general, we observed more overlap between up-regulated DEGs than down-regulated DEGs among the three genetic backgrounds.

**Figure 1.**
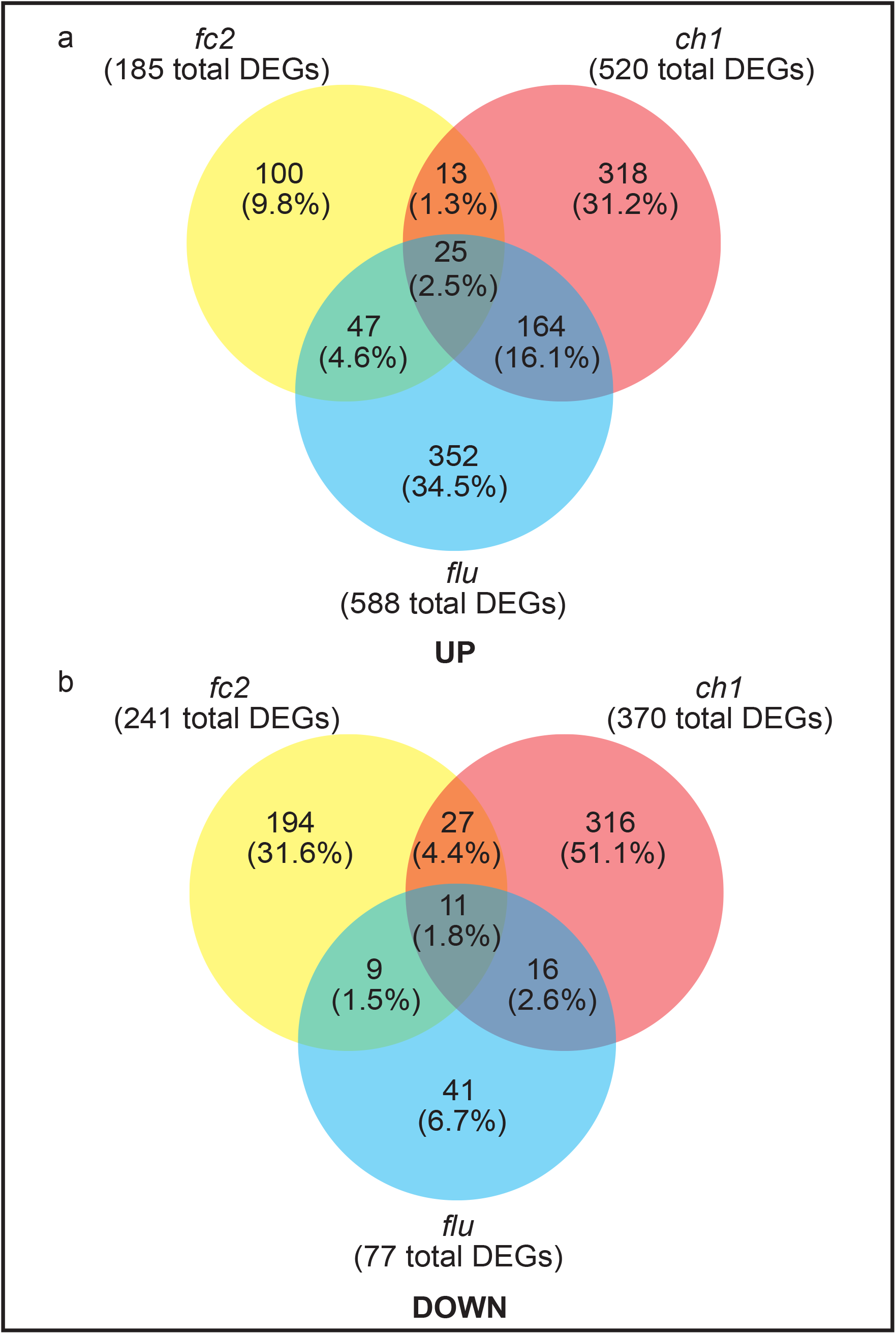
Meta-analysis of transcriptome expression data from three chloroplast ^1^O_2_-producing mutant backgrounds. Transcriptome datasets from three *Arabidopsis thaliana* ^1^O_2_-producing mutants were analyzed; *fc2* (Woodson et al. 2015), *flu* (op den Camp et al. 2003), and *ch1* (Ramel et al. 2013). Shown are Venn diagrams of differentially expressed genes (DEGs) **A)** up-regulated or **B)** down-regulated during chloroplast ^1^O_2_ stress in *fc2, flu*, and *ch1*. A ≥ 2-fold change and ≤ 0.05 adjusted p-value cutoffs were applied for all datasets where applicable. Additional information on the experiments are provided in Table S16. Tables S11 and 12 list the DEGs. Within each circle are the number of shared DEGs and the percentage of total DEGs (1,633) from the analysis.

Furthermore, we compared these DEGs to those identified in β-cc–treated wt plants (Fig S1A, B and Table S18). While we observed overlap with each mutant (8.0-18.7%), the largest overlap was observed between β-cc– treated wt plants and EL treated *ch1* mutants (18.7%). ESORGs identified in *flu* seedlings also showed a degree of overlap with each mutant (4.8-11.7%), but the largest overlap was observed with adult *flu* mutants (11.7%) (Fig. S1C and Table S19). Together, these results suggest that a core transcriptional response to chloroplast ^1^O_2_ occurs regardless of stress type, life stage, or stress duration.

To uncover roles of the overlapping gene expression, we conducted Gene Ontology (GO) term enrichment analyses. For genes upregulated in at least two genetic backgrounds (*fc2, flu*, or *ch1*), we found a diverse array of GO terms (Fig. S2A). However, the majority clustered around “response to stress,” “regulation of cellular processes,” and “aromatic compound biosynthetic process.” For genes down-regulated in two or more genetic backgrounds (Fig. S2B), we identified fewer GO terms and they had lower significance scores. Nonetheless, we observed four terms associated with photosynthesis: “pigment metabolic process,” porphyrin-containing compound metabolic process,” “tetrapyrrole metabolic process,” and “photosynthesis, light harvesting in photosystem I.” Our result is consistent with earlier studies indicating ^1^O_2_ signals reduce the expression of photosynthesis protein-encoding genes to minimize photo-oxidative damage in the light (Page et al. 2017).

Continuing our GO term enrichment analysis, we tested the DEGs in common with β-cc-treated wt and at least one other genetic background (*fc2, flu*, or *ch1*). For up-regulated genes, we observed an enrichment for many GO terms found in common between the genetic backgrounds (Fig. S3A), with one large cluster around “response to chemical.” Some small differences include GO terms related to “sulfur compound metabolic process” and “plant organ senescence.” Despite these differences, the overall similarity suggests β-cc induces a similar response to the genetic backgrounds under photo-oxidative stress, further implicating this secondary metabolite in photo-oxidative stress signaling. As before, we found fewer GO terms with less significance with the down-regulated genes, yet we identified three terms associated with cell wall modifications (Fig. S3B). These results indicate plants may change their cell walls during ^1^O_2_ stress.

Finally, we performed a GO term enrichment analysis of ESORGs (Dogra et al. 2017) up-regulated in at least one genetic background (Fig. S4). We observed a striking similarity with GO terms identified through comparing mutants (Fig. S2A), having clusters around “response to stress,” “regulation of response to stress,” and “aromatic compound biosynthetic process.” We partly expected this result as Dogra et al. (2017) identified these ESORGs from *flu* seedlings. However, we also observed an enrichment for the GO terms “cellular response to hypoxia” and “cellular ketone metabolic process,” the latter suggesting a role for secondary metabolite synthesis or signaling.

Overall, the similarity between the GO term analyses of up-regulated genes within the datasets suggest plants have a core transcriptional response to ^1^O_2_ stress to induce the expression of genes broadly involved with stress, signaling, and secondary metabolites. However, our analysis shows a large number of unique DEGs attributed to each mutant and condition suggesting that plants use different pathways depending on the specific source and site of chloroplast ^1^O_2_ stress. To delve deeper into the uniqueness of each genotype’s response to chloroplast stress, we identified the top 28 significant GO terms associated with DEGs unique to each background totaling 61 different GO terms (Table S20). We did not include down-regulated genes in this analysis as the *flu* dataset contained 41 genes, too few for a robust enrichment analysis. Each mutant had GO terms unique to itself (64%, 50%, and 43% of the GO terms for *fc2, flu*, and *ch1*, respectively). For the *fc2* mutant, GO terms involving heat and hypoxia were unique including “response to heat” “response to hypoxia” and “response to decreased oxygen levels.” For the *flu* mutant, GO terms involving defense were unique including “response to bacterium,” “defense response to bacterium,” and “regulation of defense response.” The *ch1* mutant had the fewest unique GO terms, but they included “response to hormone,” “response to abscisic acid,” and “response to jasmonic acid.” Despite these differences, we found a 28% overlap of the GO terms present in at least two mutants and a 10% overlap among all three mutants (notable GO terms include “response to stimulus,” “response to stress,” and “response to chemical.”) These results illustrate these mutant backgrounds activate similar responses (as indicated by shared and related GO terms), but utilize unique gene sets for tailored responses.

### Testing genetic interactions with the *fc2* signaling pathway in seedlings

Because the *fc2, flu*, and *ch1* ^1^O_2_-producing backgrounds all conditionally trigger to cell death and have overlapping nuclear responses (Fig. 1), we tested if they employ the same mechanisms to transmit chloroplast stress signals. Therefore, we tested if genetic suppressors identified for one ^1^O_2_-producing mutant would suppress the others. First, we introduced the *ppr30, cry1, oxi1, pub4, ex1*, and *ex2* mutant alleles into the *fc2* background. We previously demonstrated that *ex1* alone could not suppress cell death or nuclear signaling in the *fc2* mutant ((Woodson et al. 2015) and repeated those results here (Fig. S5A-C)). Because EX1 and EX2 may have partially redundant functions (Page et al. 2017), we also introduced two alleles of *ex2* (*Salk_021694/ex2-2* and *Salk_121009/ex2-3* with T-DNA insertions in the eighth exon and tenth intron, respectively) (Fig. S6A). Researchers previously showed *ex2-2* is a null allele (Uberegui et al. 2015), and our analysis confirmed this conclusion. A semi-quantitative analysis of *EX2* transcripts showed *ex2-2* is likely a null allele due to the inability to detect full-length transcript (Fig. S6B). On the other hand, *ex2-3* produced normal length transcripts and a sequencing analysis of the amplified *ex2-3* cDNA revealed normal splicing across the tenth intron. Furthermore, a RT-qPCR analysis showed wt levels of *EX2* transcript in the *ex2-3* mutant (Fig. S6C). As such, we continued our analysis with the *ex2-2* null allele.

When grown under constant 24 h light, *fc2* mutant seedlings appear pale, but healthy (Fig. 2A). However, when they grow under 6 h light / 18 h dark diurnal cycling light conditions, the seedlings bleach and die, whereas wt is unaffected. As expected, the *ppr30-1* and *pub4-6* mutations suppress the bleaching phenotype and keep the seedlings green and alive. However, we did not observe any suppression of bleaching by *cry1-304, oxi1*, or the *ex1 ex2-2* allele combination. To confirm these phenotypes, we stained the seedlings with trypan blue to assess cell death in cotyledons. As expected, *fc2* mutants stained dark blue after growing under 6 h light / 18 h dark diurnal cycling light conditions, confirming cell death (Fig. 2B and C). As expected from the visual phenotypes, *ppr30-1* and *pub4-6* significantly reduced cell death in *fc2*, while *cry1-304* and *oxi1* did not. Surprisingly, the *fc2 ex1 ex2-2* mutant did not suffer significant levels of cell death despite having a bleached appearance.

**Figure 2.**
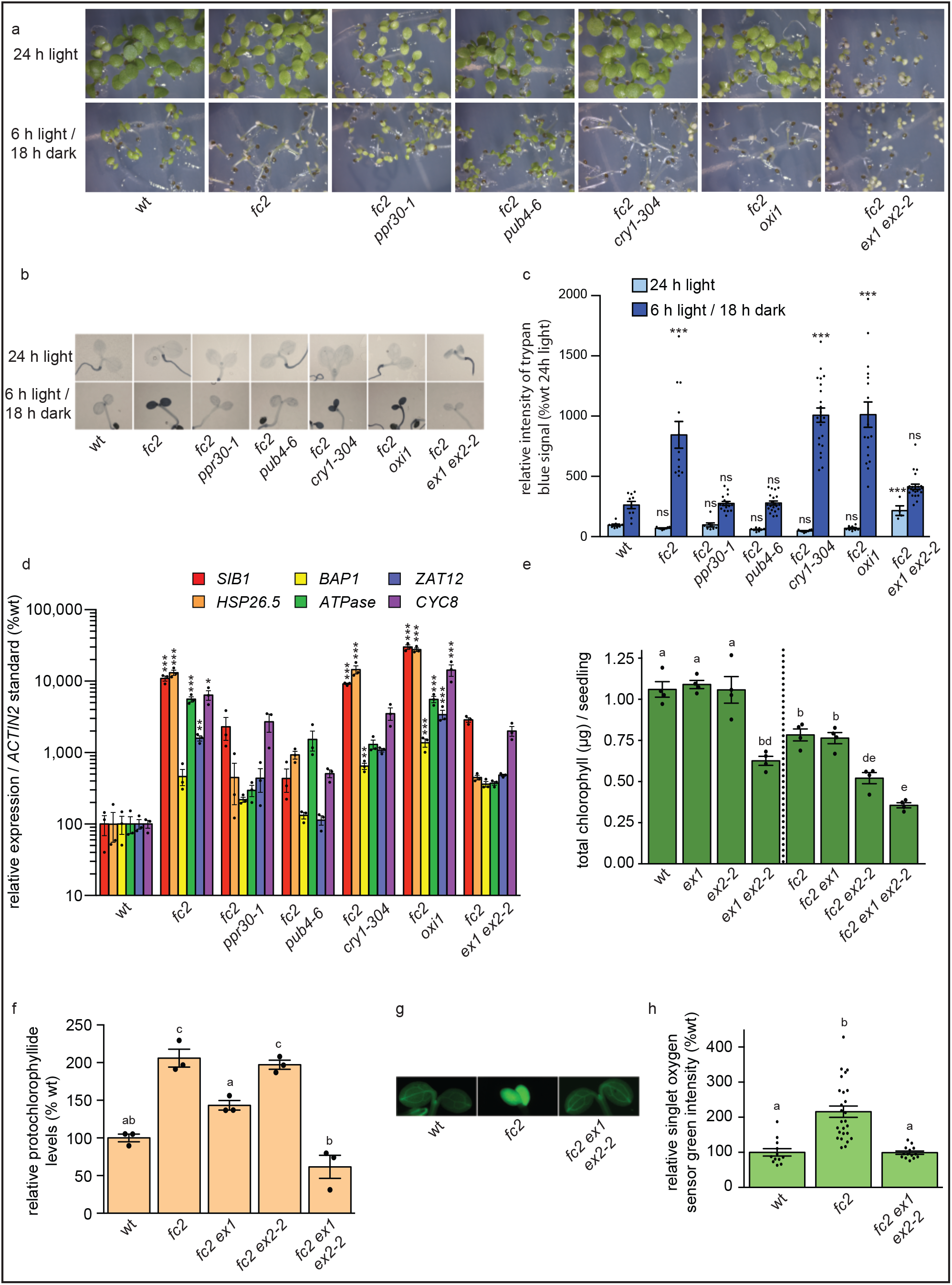
Genetic analysis of singlet oxygen signaling in *fc2* mutant seedlings. Genetic suppressors of ^1^O_2_-producing mutants were tested for their ability to suppress the stress phenotypes of *fc2* seedlings. **A)** Shown are six-day-old seedlings grown under constant light (24 h) or diurnal cycling light (6 h light / 18 h dark) conditions. **B)** Shown are representative trypan blue stains of these seedlings. The dark blue color is indicative of cell death. **C)** Shown are mean intensities of trypan blue (+/-SE, n ≥ 4 seedlings) from **B. D)** RT-qPCR analysis of stress gene markers of four-day-old seedlings grown under 6 h light / 18 h dark conditions harvested 1 h after dawn. Shown are mean expression values (+/-SE, n = 3 biological replicates). Statistical analyses in **C** and **D** were performed using a one-way ANOVA followed by a Tukey HSD test. Statistical significance in respect to wt is indicated as follows: n.s. = p-value ≥ 0.05, * = p-value ≤ 0.05, ** = p-value ≤ 0.01, *** = p-value ≤ 0.001. **E)** Mean levels of total chlorophyll (per seedling) of seven-day-old seedlings grown in 24 h light (+/-SE, n = 3 replicates). **F)** Mean levels of protochlorophyllide (Pchlide) of four-day-old dark grown (etiolated) seedlings (+/-SE, n = 3 replicates). **G)** Shown are representative images of four-day-old seedlings stained with Singlet Oxygen Sensor Green (SOSG). Seedlings were grown for three days in 6 h light / 18 h dark diurnal light cycling conditions, dark incubated at the end of day three, and re-exposed to light on day four. Pictures were acquired 3 h post-dawn. **H)** Shown are mean SOSG intensities (+/-SE, n ≥ 12 seedlings) of these seedlings. Statistical analyses in **E, F**, and **H** were performed using a one-way ANOVA followed by a Tukey HSD test. Different letters indicate statistical differences (p ≤ 0.05). In bar graphs, closed circles represent individual data points.

Next, we tested if these mutations affect retrograde signaling to the nucleus and alter the transcriptional response. We measured steady state transcript levels in four-day-old seedlings grown under 6 h light / 18 h dark diurnal cycling light conditions one hour post subjective dawn using RT-qPCR, probing for six previously identified chloroplast stress marker genes (*SIB1* and *HSP26*.*5* identified in ^1^O_2_-stressed *fc2* seedlings (Woodson et al. 2015), *BAP1* and *ATPase* identified in ^1^O_2_-stressed *flu* seedlings (op den Camp et al. 2003), and general oxidative stress markers *ZAT12* and *GST* (Baruah et al. 2009)). As shown in Fig. 2D, photo-oxidative stress significantly induces expression of five of the six stress marker genes (excluding the *flu* marker *BAP1*) in *fc2* compared to wt. As expected of suppressors, both *ppr30-1* and *pub4-6* reduce induction of these marker genes. In line with their bleached phenotypes, *cry1-304* and *oxi1* did not hugely impact of expression of the marker genes. Compared to wt, *fc2 cry1-304* and *fc2 oxi1* experienced significant induction of all marker genes (except for *ZAT12* in *fc2 cry1-304*). Despite its pale appearance, the *fc2 ex1 ex2-2* mutant transcriptionally resembled the suppressors (*fc2 ppr30-1* and *fc2 pub4-6*) with no significant induction of stress marker genes compared to wt. Together, these results suggest neither CRY1 nor OXI1 play a major role in ^1^O_2_-triggered cell death or retrograde signaling in *fc2* mutant seedlings. However, the results reveal a potential genetic interaction between *fc2* and the *ex1 ex2-2* combination.

We did not expect the *ex1 ex2-2* combination to suppress cell death and transcriptomic responses in *fc2* as *ex1* does not partially suppress these *fc2* phenotypes alone ((Woodson et al. 2015) and Fig. S5A-C). To distinguish if *ex1* and *ex2-2* additively suppress cell death or if *ex2-2* alone is sufficient, we generated an *fc2 ex2-2* mutant. Under 6 h light / 18 h dark diurnal cycling light conditions, the *fc2 ex2-2* mutant was visually similar to the *fc2* mutant (Fig. S5A). Furthermore, trypan blue stains confirmed the *ex2-2* mutation alone did not suppress cell death in the *fc2* background (Figs. S5B and C).

One possible mechanism to suppress cell death in *fc2* is through reducing tetrapyrrole biosynthesis (either directly or by reducing general chloroplast development). Second site mutations can accomplish this reduction by decreasing flux through the tetrapyrrole pathway and avoiding ^1^O_2_ accumulation (e.g. plastid protein import and tetrapyrrole biosynthesis mutants) (Woodson et al. 2015). Indeed, the *fc2 ex1 ex2-2* mutant appeared very pale even under permissive 24 h constant light conditions (Fig. 2A). As expected, these mutant seedlings had significantly reduced levels of total chlorophyll compared to *fc2* (Fig. 2E). *ex1* and *ex2-2* had an additive effect in terms of chlorophyll reduction (the triple mutant contained less total chlorophyll than either *fc2 ex* double mutant) independent of the *fc2* background (*ex1 ex2-2* accumulated less total chlorophyll than wt). To determine if light-induced degradation or decreased tetrapyrrole synthesis caused a reduction in chlorophyll levels, we measured steady-state protochlorophyllide (Pchlide) levels in etiolated (dark grown) seedling to gauge the flux through the tetrapyrrole pathway. As previously shown, *fc2* mutants accumulate two-to-three-fold excess Pchlide compared to wt ((Woodson et al. 2015) and Fig. 2F). The *fc2 ex1 ex2-2* mutant had wt Pchlide levels suggesting the mutant had reduced tetrapyrrole synthesis. Next, we measured bulk ^1^O_2_ levels in four-day-old seedlings grown under 6 h light / 18 h dark diurnal cycling light conditions using Singlet Oxygen Sensor Green (SOSG) (Fig. 2G and H). Two hours after subjective dawn, *fc2* mutants accumulated excess ^1^O_2_ compared to wt. However, the *fc2 ex1 ex2-2* mutant had wt ^1^O_2_ levels. Together, these results suggest the *ex1* and *ex2-2* mutations additively block *fc2* phenotypes by reducing tetrapyrrole synthesis and ^1^O_2_ production rather than by directly affecting a signaling mechanism as shown in the *flu* mutant.

### Testing genetic interactions with the *fc2* signaling pathway in adult plants

As life stage could affect the ability of these mutations to suppress the *fc2* cell death phenotype, we tested for suppression of cell death in adult plants. We grew plants for 21 days under 24 h constant light conditions and shifted them to 16 h light / 8 h dark diurnal cycling light conditions for 6 days. As a control, we kept another set of plants in 24 h constant light for the full 27 days. Under constant 24 h light, *fc2* plants appeared relatively healthy and do not present any indications of obvious cell death lesions (Fig. 3A). However, after shifting to 16 h light / 8 h dark diurnal cycling light conditions, *fc2* mutants developed leaf lesions. A trypan blue stain confirmed these lesions are areas of cell death (Figs. 3B and C). If a mutation causes suppression of ^1^O_2_ signaling in *fc2*, we expect a reduction in the appearance of leaf lesions under these conditions. We found, as expected, that the *ppr30* and *pub4-6* mutations suppressed the *fc2* cell death phenotype, having fewer observable lesions and less trypan blue staining than the *fc2* single mutant (Figs. 3A-C). Surprisingly, we found the *oxi1* mutation significantly suppressed lesion formation in the *fc2* mutant, suggesting OXI1 may play a role in ^1^O_2_ signaling in *fc2* adult plants. The *ex1 ex2-2* combination did not suppress cell death in adult leaves, consistent with these mutations leading to developmental (rather than signaling) defects. As in seedlings, *cry1-304* did not suppress cell death, further suggesting CRY1 does not play a strong role in ^1^O_2_ signaling in the *fc2* mutant.

**Figure 3.**
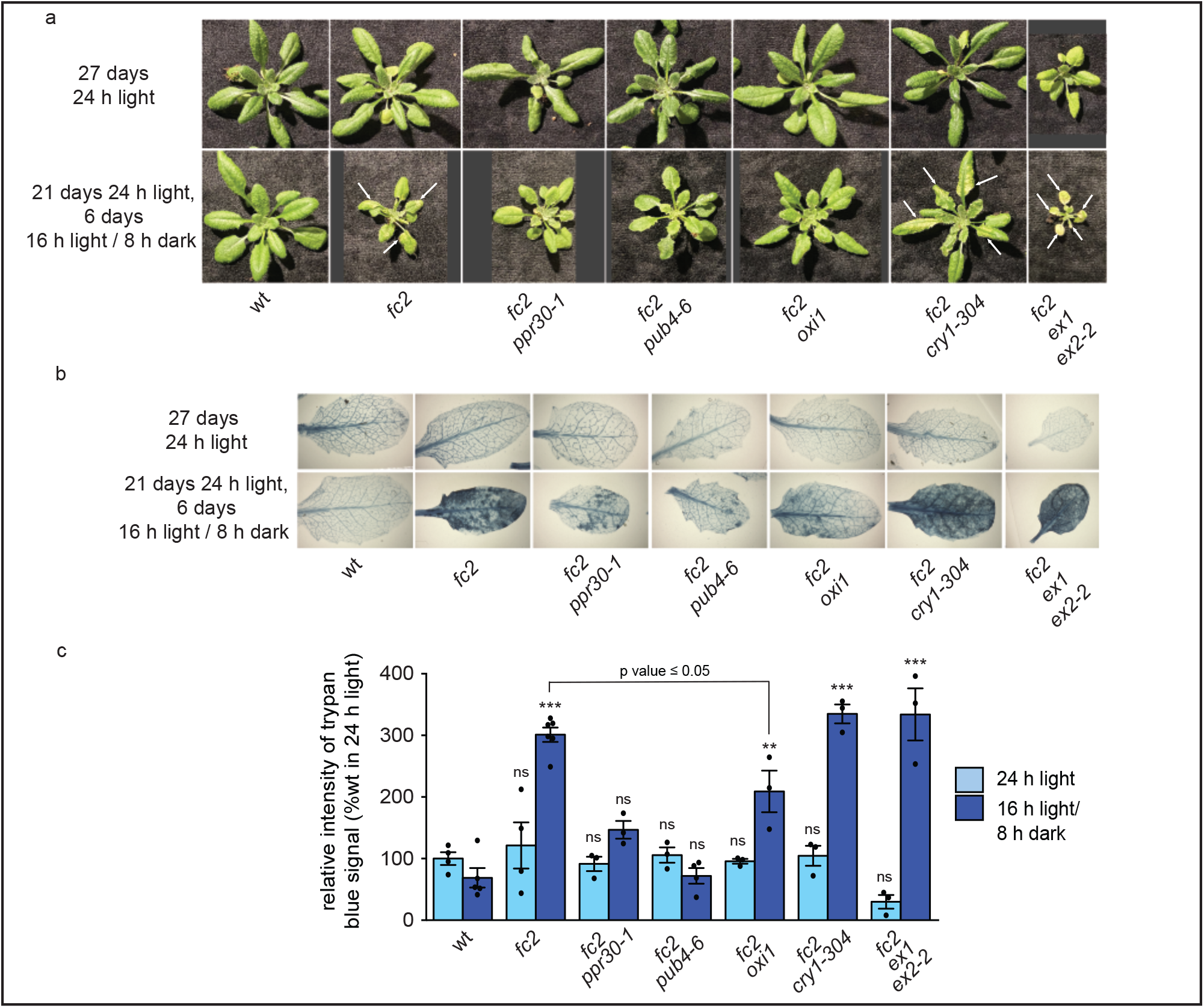
Genetic analysis of singlet oxygen signaling in *fc2* mutant adult plants. Genetic suppressors of ^1^O_2_-signaling were tested for their ability to suppress the cell death phenotype of *fc2* adult plants. **A)** Shown are representative 27-day-old plants grown under constant light (24 h) or under 24 h light for 21 days and then shifted to diurnal cycling light conditions (16 h light / 8 h dark) for 6 additional days. White arrows indicate lesions. **B)** Shown are representative trypan blue cell death stains for both sets of plants. The dark blue color is indicative of cell death. **C)** Shown are mean intensities of trypan blue (+/-SE, n ≥ 3 leaves) from **B**. Statistical analyses within each light treatment were performed using a one-way ANOVA followed by a Tukey HSD test. Statistical significance in respect to wt is indicated as follows: ** = p-value ≤ 0.01, *** = p-value ≤ 0.001, not significant (ns) = p-value ≥ 0.05. Closed circles represent individual data points.

### Testing genetic interactions in *flu* mutant seedlings

To continue our assessment of potential genetic interactions of known chloroplast ^1^O_2_ suppressors in other ^1^O_2_-generating backgrounds, we monitored phenotypes of ^1^O_2_ signaling mutations in the *flu* mutant background. Previously, researchers determined *flu* mutants accumulate chloroplast ^1^O_2_ proportionally to increasing lengths of time in the dark (Wang et al. 2020). To experimentally determine the length of time in the dark needed to induce cell death, we treated five-day-old wt and *flu* seedlings to various lengths of time in the dark (0, 4, 8, 12, and 24 h) and re-exposed the seedlings to light for 36 h. Based on the outcomes shown in Fig. S7A, we decided 12 h of dark was adequate to completely bleach most *flu* seedlings within 36 h of light exposure. Therefore, we crossed in ^1^O_2_ signaling mutations (*ex1, pub4-6, cry1-304*, and *oxi1*) into the *flu* background to test which mutations may block ^1^O_2_ signaling phenotypes.

As expected, the *ex1* mutation suppressed bleaching in the *flu* seedlings after a 12 h dark treatment (Fig. 4A), while the other mutations did not obviously appear to affect bleaching. To confirm our visual assessment, we performed a trypan blue stain using the 12 h time point to confirm *flu* mutants experience extensive cell death in their cotyledons, and the *ex1* mutation significantly reduces this effect to near wt levels (Figs. 4B and C). As expected from their bleached phenotypes, *flu pub4-6* and *flu oxi* stained similarly to *flu*. However, we found *flu cry1-304* stained significantly lower than *flu* (p value ≤ 0.001), confirming CRY1 plays at least a minor role in ^1^O_2_ signaling in *flu* mutants (Danon et al. 2006).

**Figure 4.**
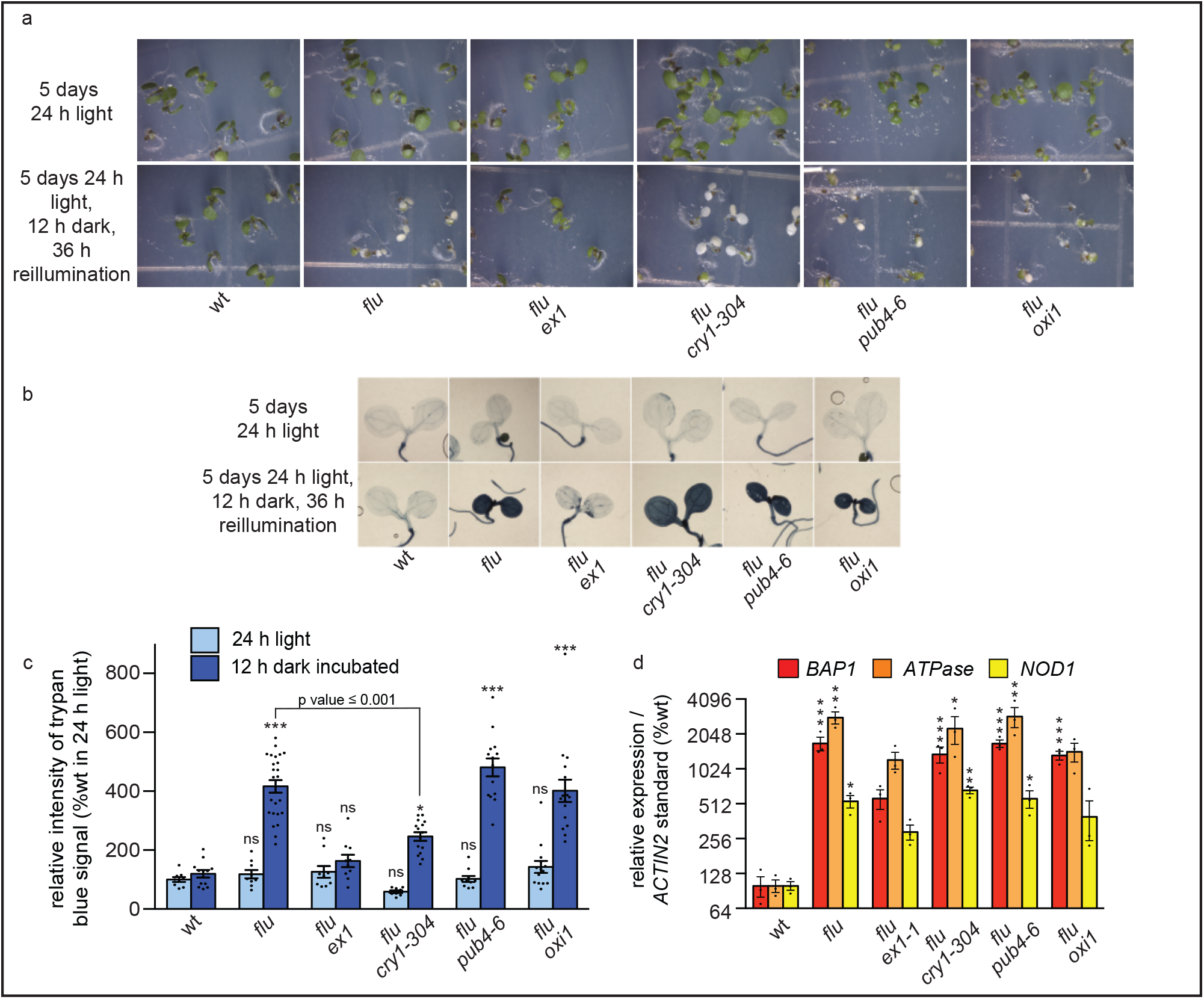
Genetic analysis of singlet oxygen signaling in *flu* seedlings. Genetic suppressors of chloroplast ^1^O_2_ signaling were tested for their ability to suppress *flu* phenotypes in seedlings. **A)** Shown are (top) six-day-old seedlings grown under constant light (24 h) and (bottom) five-day-old seedlings grown under 24 h light, incubated in the dark for 12 h, and re-exposed to light for 36 h (7 days total). **B)** Shown are representative images of these seedlings stained with trypan blue. The dark blue color is indicative of cell death. **C)** Mean intensities of trypan blue signal (+/-SE, n ≥ 9 seedlings) from panel **B. D)** RT-qPCR of *flu*-specific stress gene markers of five-day-old seedlings grown under 24 h constant light then dark-incubated for 12 h, harvested 1 h after re-exposure to light. Shown are mean expression values (+/-SE, n ≥ 3 biological replicates). Statistical analyses were performed using a one-way ANOVA followed by a Tukey HSD test. Statistical significance in respect to wt is indicated as follows: n.s. = p-value ≥ 0.05, * = p-value ≤ 0.05, ** = p-value ≤ 0.01, *** = p-value ≤ 0.001. Closed circles represent individual data points.

Next, we tested if our double mutants activated ^1^O_2_–triggered retrograde signaling by measuring the expression of stress marker genes (as for *fc2* (Fig. 2D)). Here, we also included *NOD1*, another gene induced in *flu* mutants (op den Camp et al. 2003). We placed five-day-old seedlings in the dark for 12 h and exposed them to 1 h light prior to tissue collection for RNA extraction. In the *flu* mutant, we observed significant induction of all three *flu*-specific stress marker gene transcripts (Figs. 4D). Expectedly, we observed that *ex1* reduced expression of these genes (Lee et al. 2007). *pub4-6* and *cry1-304* did not significantly reduce any one of these transcripts. However, *oxi1* lowered levels of two transcripts (*ATPase* and *NOD1*). We tested the other four stress marker transcript levels in the *flu* mutant. We observed higher transcript levels compared to wt, but they were not significant (Fig. S7B). Furthermore, none of the suppressor mutations significantly reduced expression of these marker genes. Together, these results suggest PUB4 does not play a significant role in facilitating the ^1^O_2_ signal in *flu* mutants, while CRY1 plays a minor role in regulating cell death in *flu* seedlings. OXI1 does not play a major role in triggering cell death in the *flu* mutant, yet it may play a minor role in transmitting the retrograde signal to the nucleus.

### Testing genetic interactions in *flu* adult plants

As with the *fc2* mutant, we assessed if life stage affected ^1^O_2_-signaling in the *flu* mutant. We grew wt, *flu*, and the double mutant plants under 24 h constant light conditions to avoid ^1^O_2_ stress. We then shifted them to 16 h light / 8 h dark diurnal cycling light conditions for 3 days to accumulate Pchlide and ^1^O_2_. As expected, the *flu* plants developed lesions under these conditions, whereas wt appeared normal (Fig. 5A). *flu ex1* plants exposed to 16 h light / 8 h dark diurnal cycling light conditions for 5 days did not develop leaf lesions, consistent with EX1 playing a role in ^1^O_2_-triggered cell death regardless of life stage (Wagner et al. 2004) (Fig. 5A). As in seedlings, we did not observe obvious suppression of the cell death phenotype by *cry1-304, pub4-6*, or *oxi1* in adult plants. We confirmed these cell death phenotypes with a trypan blue cell death stain (Figs. 5B and C). Together, we conclude ^1^O_2_ signaling in the *flu* mutant utilizes the *EX1*-dependent pathway rather than the PUB4 (and possibly OXI1)-dependent chloroplast quality control pathway implemented by *fc2*. However, OXI1 may contribute to the retrograde signaling in seedlings to control some nuclear gene expression in the *flu* mutant.

**Figure 5.**
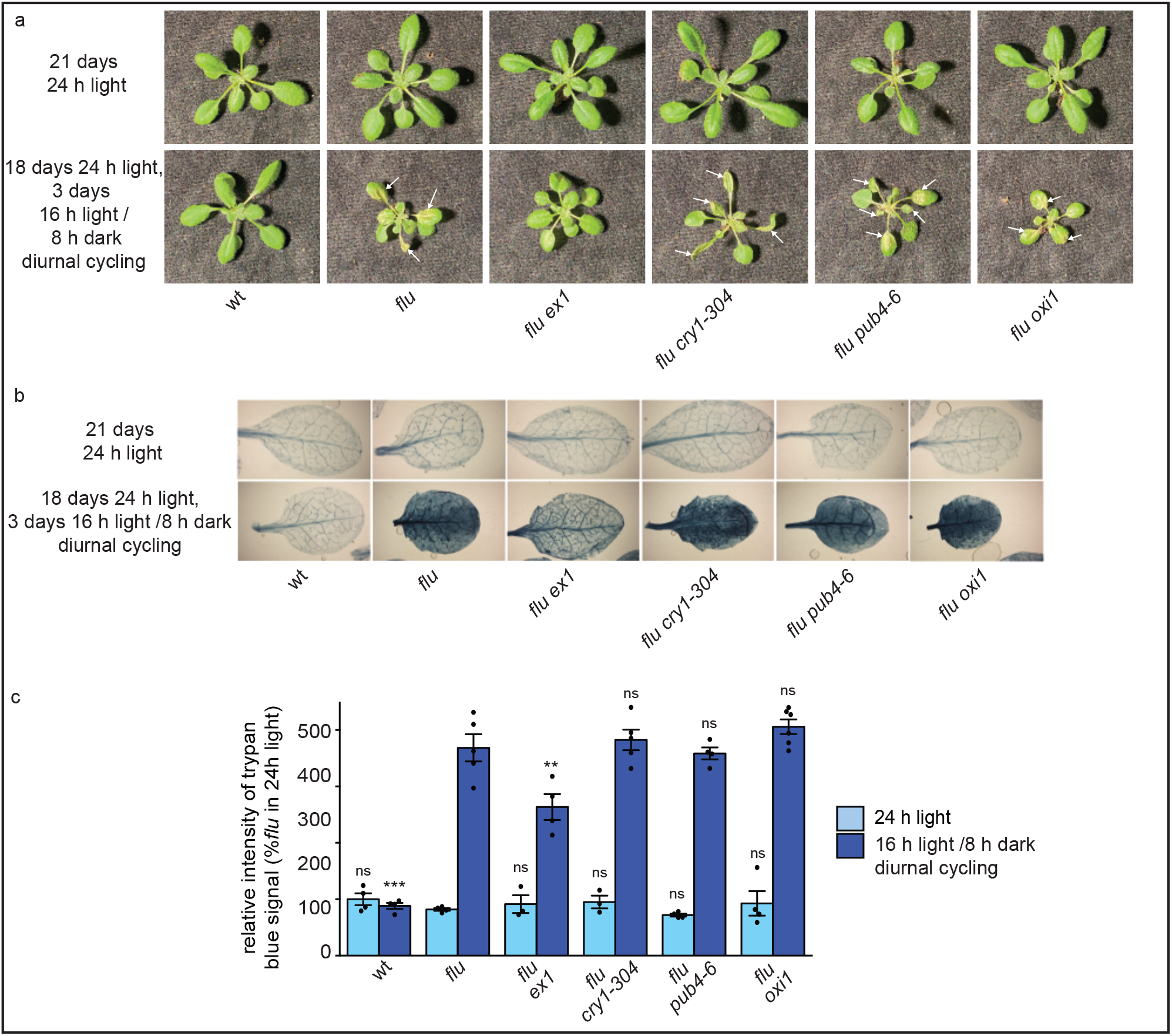
Genetic analysis of ^1^O_2_ signaling in adult *flu* plants Genetic suppressors of ^1^O_2_-signaling were tested for their ability to suppress the *flu* cell death phenotype in adult plants. **A)** Shown are representative 24-day-old plants grown under constant light (24 h) and 21-day-old plants grown in 24 h light and then shifted to diurnal cycling light (16 h light / 8 h dark) conditions for three days (24 days old total). White arrows indicate lesions. **B)** Shown are representative images of leaves from these plants stained with trypan blue. The dark blue color is indicative of cell death. **C)** Shown are mean intensities of trypan blue (+/-SE, n ≥ 3 leaves) from **B**. Statistical analyses within each light treatment were performed using a one-way ANOVA followed by a Tukey HSD test. Statistical significance in respect to *flu* is indicated as follows: ** = p-value ≤ 0.01, *** = p-value ≤ 0.001, not significant (ns) = p-value ≥ 0.05. Closed circles represent individual data points.

### Testing genetic interactions with the *chlorina1* signaling pathway

Previously, researchers demonstrated that growing the *ch1* mutant under EL stress (≥ 1,100 μmol photons m^-2^ sec^-1^) induces ^1^O_2_ signaling (Ramel et al. 2013). The generated ^1^O_2_ initiates cell death, which the *oxi1* mutation blocks in adult plants (Shumbe et al. 2016). Furthermore, researchers demonstrated EX1 and EX2 are not involved in *ch1*’s ^1^O_2_ signaling since *ch1 ex1 ex2* mutants suffer from a comparable level of EL-triggered cell death to *ch1* (Ramel et al. 2013). Here, we tested the involvement of PUB4 in transmitting this ^1^O_2_ signal. We grew seedlings under permissive light conditions (120 μmol photons m^-2^ sec^-1^) for 7 days and shifted them to 1,200 μmol photons m^-2^ sec^-1^ for 24 h. We lowered the ambient temperature to 10°C to avoid any incidental heat stress caused by the increased radiation. Within 2 h, all seedlings experienced a decrease in maximum photosystem II quantum efficiency (F_v_/F_m_), which continued to decrease for 24 h of treatment (Fig. 6A). Furthermore, we observed photo-bleaching of cotyledons after 12 h of EL. Photo-bleaching worsened after 24 h (Fig. 6B). At the same time points, we observed increased susceptibility (lowered F_v_/F_m_ values and increased bleaching) of the *ch1* mutant to EL stress, consistent with its chloroplasts experiencing increased photo-damage (Ramel et al. 2013). The *pub4-6* mutation partially reversed these effects (increased F_v_/F_m_ values at 2 and 6 h EL and delayed bleaching at 12 h EL), while the *oxi1* mutation did not reverse them. Additionally, we observed increased tolerance of the *pub4-6* single mutant to EL compared to wt, having higher F_v_/F_m_ values at 2 and 6 h EL and delayed bleaching at 12 h EL (Figs. 6A and B). Together, these results suggest PUB4 may be involved in EL-triggered ^1^O_2_ signaling in the seedling stage.

**Figure 6.**
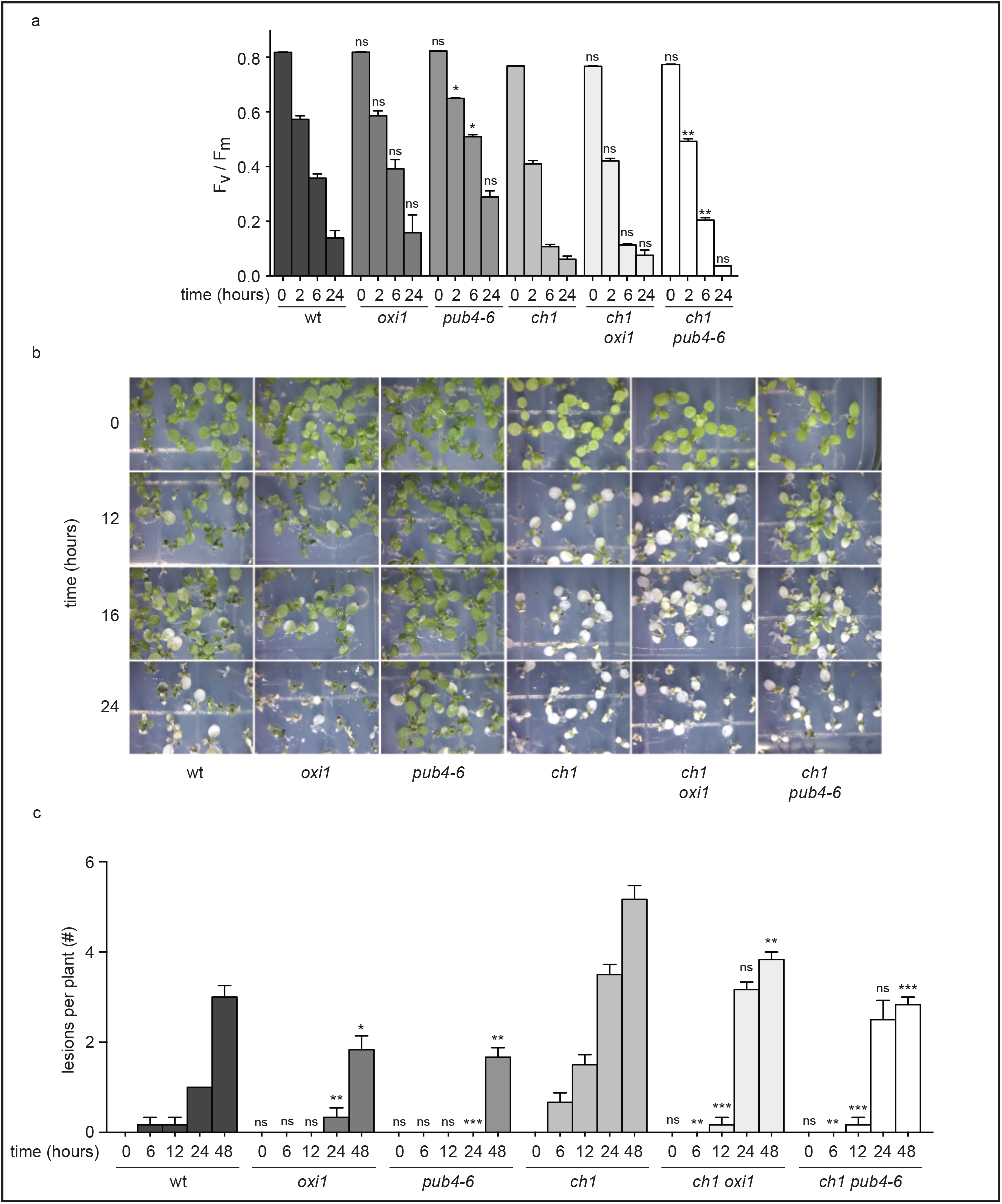
Effect of singlet oxygen signaling mutations on excess light-induced phenotypes. The effect of the *oxi1* and *pub4-6* mutations were tested in excess light (EL) conditions. **A)** Time course analysis of maximum PSII quantum efficiency (F_v_/F_m_) in seven-day-old seedlings during 24 h of EL (1,200 μmol photons sec^-1^ m^-2^) at 10°C (+/-SE, n > 3 groups of seedlings). **B)** Shown are representative seedlings immediately after the indicated length of EL treatment. **C)** Time course analysis of F_v_/F_m_ in 18-day-old adult plants during 48 h of EL (1,300 μmol photons sec^-1^ m^-2^ (+/-SE, n ≥ 6 plants). Statistical analyses within each time point (for wt or *ch1*) were performed using a one-way ANOVA followed by a Tukey HSD test. Statistical significance in respect to wt (for *oxi1* and *pub4-6*) or *ch1* (for *ch1 oxi1* and *ch1 pub4-6*) is indicated as follows: * = p-value ≤ 0.05, ** = p-value ≤ 0.01, *** = p-value ≤ 0.001, not significant (ns) = p-value ≥ 0.05.

As researchers previously studied the *ch1* EL-treated phenotype in adult plants and leaves (Ramel et al. 2013; Shumbe et al. 2016), we tested EL sensitivity in 18-day-old plants. We grew plants under permissive light conditions (70 μmol photons m^-2^ sec^-1^) for 18 days. Next, we shifted them to 1,300 μmol photons m^-2^ sec^-1^ at 10°C. Similar to seedlings, all plants experienced an immediate decrease in F_v_/F_m_ values indicating photo-damage (Fig. S8A). Again, the *ch1* mutant was particularly susceptible. *oxi1* and *pub4-6* did not significantly affect F_v_/F_m_ values in the *ch1* background (although *pub4-6* single mutants had significantly higher F_v_/F_m_ values compare to wt at 6 and 12 h EL). However, *oxi1* and *pub4-*6 attenuated photo-bleaching in both wt and *ch1* backgrounds (Figs. 6C and S8B). Together, our results suggest both OXI1 and PUB4 play a role in transmitting EL-triggered stress signals, but *OXI1* may play a stage-specific role in adult leaves.

### Onset of lesions in the *acd2* mutant was slowed by *pub4-6*

As *pub4-6* mitigated ^1^O_2_-induced cell death in the *fc2* and *ch1* mutants, we investigated if it can affect lesion formation caused by other sources of ROS. Therefore, we examined the effect of the *pub4-6* mutation in the ROS and lesion accumulating *acd2* mutant. Prior work determined that a *cry1* mutation or a *ex1 ex2* combination did not significantly alter the accumulation of lesions, suggesting the *flu* signaling pathway is not being used in *acd2* to induce cell death (Pattanayak et al. 2012). As PUB4 appears to represent another separate ^1^O_2_ pathway, we tested if PUB4 plays a role in lesion formation in *acd2* mutants by generating *acd2 pub4-6* double mutants and growing them alongside wt and the corresponding single mutants under 16 h light / 8 h dark diurnal light cycling conditions. Initially, *acd2* mutant plants appeared healthy and comparable to wt. However, after 18 days, *acd2* mutants began to randomly develop lesions of cell death on their leaves, which accumulated until senescence (Figs. 7A and B and S9). Conversely, we observed the *acd2 pub4-6* double mutant appeared healthier than the *acd2* single mutant, and developed fewer leaves with lesions over time. At 36 days, we stopped the experiment as we could not reliably distinguish between *acd2*-specific and normal leaf senescence lesions. This early senescence was particularly apparent in *pub4-6* as previously reported (Woodson et al. 2015). Together, these data suggest PUB4 is involved in regulating ROS-induced lesion formation in *acd2* plants.

**Figure 7.**
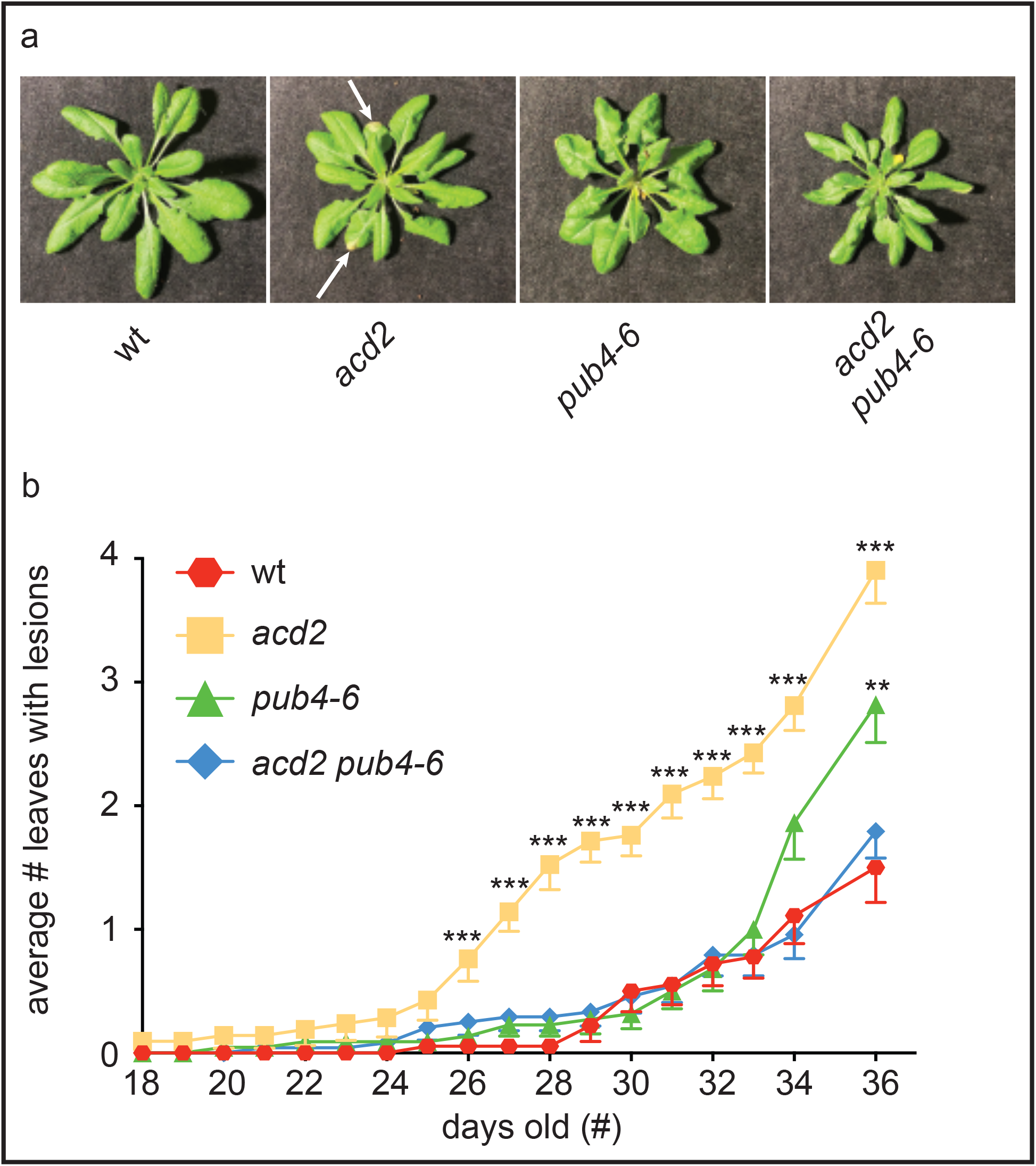
The *pub4-6* mutation slows the progression of spontaneous cell death in the *acd2-2* mutant. The *acd2* spontaneous cell death and lesion phenotypes were assessed. **A)** Shown are representative 30-day-old adult plants grown under 16 h light / 8 h dark diurnal cycling light conditions. The inflorescences were removed for the picture. White arrows indicate lesions. **B)** Mean number of leaves with lesions per plant (+/-SE, n ≥ 18 plants). Statistical analyses were performed for each time point using a one-way ANOVA followed by a Tukey HSD test. Statistical significance as follows in respect to wt: ** = p-value < 0.01, *** = p-value < 0.001.

## Discussion

Despite knowing chloroplast ^1^O_2_ has signaling capabilities, natural stresses complicate its study by causing complex ROS signatures in plant cells (Choudhury et al. 2017; Rosenwasser et al. 2013). Thus, to dissect ^1^O_2_ signals, researchers use several Arabidopsis mutants that conditionally and specifically produce this ROS, including *fc2* (Woodson et al. 2015), *flu* (Meskauskiene et al. 2001), *ch1* (Ramel et al. 2013), and *acd2* (Pruzinská et al. 2007). To assess how similar these signaling pathways are, we compared publicly available transcriptomic data for the *fc2, flu*, and *ch1* mutants (the *acd2* mutant did not have an available dataset for comparison). We found each mutant shared a proportion of their DEGs with the other two mutants (Fig. 1). We were surprised by this overlap since the datasets represent samples collected from different ages (seedling vs. adult) and grown in different conditions to elicit ^1^O_2_ stress in the chloroplasts. Therefore, we hypothesize these genotypes share a core transcriptomic response and that the different sources of ^1^O_2_ converge on some transcriptomic responses. To further support our hypothesis, we found ß-cc responsive genes and ESORGs in the mutant datasets (Figs. S1A-C). Notably, as we found ß-cc responsive genes in the *fc2* and *flu* datasets, we can hypothesize these mutants produce ß-cc. To the best of our knowledge, however, researchers have not reported such measurements.

Notwithstanding the similarities found in the transcriptomic meta-analysis, we identified DEGs specific to each background (Fig. 1A). Although these DEGs were unique, we found they produced similar GO-terms between mutants for up-regulated genes (Table S20). These GO terms include “response to stimulus,” “response to stress,” and “response to chemical.” These results may indicate these mutants employ different sets of genes with similar functions for the cell to respond or acclimate appropriately to a specific stress. We hypothesize such ROS signatures may mimic those caused by different types of environmental stress as each of these mutants produce ^1^O_2_ differently (Rosenwasser et al. 2013).

Our meta-analysis revealed the potential for the existence of multiple ^1^O_2_ signaling pathways in these mutants. Therefore, we took a genetic approach to identify potential converging points of these pathways. We hypothesized that if a secondary suppressor mutation for one mutant background has a genetic interaction within a different mutant background, these two mutant backgrounds may share a portion of their signaling cascades. First, with *fc2* mutant seedlings, we observed the *ppr30-1* and *pub4-6* mutations suppressed cell death and retrograde signaling as previously described (Alamdari et al. 2020; Woodson et al. 2015). Initially, we did not notice obvious suppression of the *fc2* cell death phenotype by suppressors of the *flu* or *ch1* backgrounds (Fig. 2A). Nonetheless, with further analysis, we found *fc2 ex1 ex2-2* seedlings had blocked cell death and retrograde signaling even though they were bleached white under permissible growing conditions (Figs. 2A-C).

These results surprised us as we previously demonstrated *ex1* alone does not block ^1^O_2_ controlled retrograde signaling or cell death in *fc2* seedlings or adults (Woodson et al. 2015). Indeed, both the *fc2 ex1* and *fc2 ex2-2* double mutants had similar levels of cell death to *fc2*, suggesting the suppression in the triple mutant was additive (Figs. S5A-C). We hypothesize the mechanism of suppression is indirect, as the *fc2 ex1 ex2-2* accumulated less bulk ^1^O_2_ compared to *fc2* (Figs. 2G and H). The observed suppression is likely due to a delay in chloroplast development and a decrease in tetrapyrrole synthesis: *ex1 ex2-2* led to a decrease in chlorophyll and Pchlide levels (Figs. 2E and F). Furthermore, the *ex1 ex2-2* combination failed to block cell death in adult *fc2* mutants, which suggests that once chloroplasts become fully developed, the *ex1 ex2-2* combination does not affect the *fc2* phenotype. How the *ex1 ex2* combination affects chloroplast development is not clear as these proteins are primarily implicated in ROS signaling. Adding to the complexity, recent work suggests these two proteins have different functions. EX2 may act as a decoy to protect EX1 from oxidation and prevent premature signaling (Dogra et al. 2022). Thus, a double mutant may have a more complicated phenotype than previously assumed. Even so, we conclude the EX proteins do not play a direct role in ^1^O_2_ signaling in the *fc2* mutant.

To our surprise, the *oxi1* mutation suppressed the cell death phenotype in *fc2* adult plants (similarly to the *ppr30-1* and *pub4-6* mutations) even though it did not affect cell death or retrograde signaling in seedlings. To date, researchers have primarily studied the role of OXI1 in ^1^O_2_ signaling in adult plants and leaves (Shumbe et al. 2016). However, some research indicates OXI1 plays a role in basal defense against oomycete pathogens in seven-day-old seedling cotyledons (Rentel et al. 2004) indicating that OXI1 is present in this tissue. This may mean that OXI1 only participates in chloroplast ^1^O_2_ signaling in true leaf mesophyll cells, which is consistent with our EL experiments where the *oxi1* mutation only mitigated cell death in adult leaves (Figs. 6A-C). Prior work indicates chloroplast physiology differs between the embryonic cotyledons in seedlings and the mesophyll cells in true leaves (Albrecht et al. 2008). Whether a different serine/threonine kinase is used in cotyledons, or there is another mechanism altogether, will require further study.

Next, we performed similar experiments in the *flu* mutant. In seedlings, we observed suppression of cell death by the *ex1* and *cry1-304* mutations, and *ex1* reduced expression of all three nuclear marker genes (Figs. 4A-D). Neither *pub4-6* or *oxi1* reduced cell death, although *oxi1* did have a minor effect on retrograde signaling. While EX1 is necessary for ^1^O_2_ signaling in *flu* mutants (Dogra et al. 2019), researchers have previously demonstrated CRY1’s involvement only in protoplasts (Danon et al. 2006). In protoplasts, the effect of the *cry1* mutation was equally strong compared to *ex1* in terms of cell death. In our study, the effect of the *cry1* mutation was noticeably weaker than *ex1* in seedlings, and *cry1* did not significantly suppress cell death in the adult phase (Figs. 5B and C). These results may be due to a difference between systems (*in vitro* protoplasts vs *in planta*). The protoplast study also used the *Landsberg erecta* ecotype with different *flu* and *cry1* alleles (this study was performed in the *Columbia* background), which may also account for some of the differences. When researchers performed a microarray analysis of ^1^O_2_-responsive transcripts in *flu* protoplasts, they found the *cry1* mutation affected only ∼3% of the DEGs in *flu* (Danon et al. 2006). When we pair this finding with our work here, we hypothesize CRY1 plays a minor role in the ^1^O_2_ retrograde signal, which can be uncoupled from cellular degradation. Overall, however, our results suggest the *flu* mutant emits a functionally unique signal that does not involve PUB4 or OXI1 to control cellular degradation.

To continue our investigation of ^1^O_2_ signaling, we next tested the *ch1* mutant that produces ^1^O_2_ due to unprotected PSII reaction centers (Ramel et al. 2013) and signals for cell death with OXI1 (Shumbe et al. 2016). Previously, researchers showed cellular degradation in this background is independent of EX1 and EX2; *ch1 ex1 ex2* mutants still suffer EL-induced lesions and PSII photo-inhibition (Ramel et al. 2013). Accordingly, we assessed if the *fc2* pathway is active in *ch1* mutants by testing the role of PUB4 in EL stress. In EL-stressed seedlings, *pub4-6* attenuated cotyledon bleaching and the reduction in F_v_/F_m_ values in the wt and *ch1* backgrounds (Figs. 6A and B). Somewhat surprisingly, the *oxi1* mutation did not affect these phenotypes, further suggesting OXI1 is not involved in ^1^O_2_ signaling in seedlings. In the adult phase, both *oxi1* and *pub4-6* attenuated EL-induced lesions in the wt and *ch1* backgrounds (Fig. 6C), confirming earlier reports on OXI1 (Shumbe et al. 2016) and suggesting *fc2* and *ch1* mutants share a ^1^O_2_ signaling pathway induced by natural EL stress.

Finally, as *pub4-6* can block ^1^O_2_-induced cell death in *fc2* and *ch1* mutants, we tested if we would observe a similar trend in the *acd2* mutant, which produces leaf lesions due to the accumulation of chlorophyll breakdown products (such as RCC) that can produce ^1^O_2_ (Pruzinská et al. 2007). Remarkably, the *acd2 pub4-6* double mutant had delayed onset of lesions compared to the *acd2* single mutant, suggesting *acd2* mutants activate the same pathway as *fc2* and *ch1* mutants (Figs. 7A and B). Previously, researchers demonstrated neither *cry1* or an *ex1 ex2* combination could reduce lesion formation in the *acd2* background (Pattanayak et al. 2012), further indicating the ^1^O_2_-signaling pathway used by *flu* mutants is distinct. This result also led the authors to conclude chloroplast ^1^O_2_ and ROS were not responsible for cell death. As ^1^O_2_ can be detected in *acd2* mitochondria, these researchers hypothesized these organelles may trigger a signal instead. Nevertheless, the observation that *pub4-6* reduces lesion formation in *acd2* opens the possibility for the involvement of chloroplast ^1^O_2_ during lesion formation in *acd2* mutants. On the other hand, we cannot rule out the possibility that PUB4 plays a role in mitochondrial ROS signaling.

Overall, our work suggests chloroplasts use at least two ^1^O_2_ signaling pathways to control cellular degradation. One pathway depends on the PUB4 protein to control cellular degradation and is employed bt *fc2, ch1*, and *acd2* mutants along with EL-stressed wt plants. At least in *fc2* and *ch1* mutants, the OXI1 protein participates in this pathway (its role in *acd2* was not tested), but is restricted to true leaves in adult tissue. *flu* mutants utilize an alternative ^1^O_2_ signaling pathway. Instead, their ^1^O_2_ signal requires EX1 and (at least in seedlings) CRY1 to initiate cell death. Retrograde signaling to the nucleus to alter the transcriptome follows a similar, yet more complex, pattern. For instance, the *cry1* and *oxi1* mutations partially reduce the expression of some stress marker genes in the *fc2* and *flu* backgrounds, respectively, despite no effect on cell death. As these effects on transcript levels were relatively mild, we hypothesize some crosstalk may exist between these two ^1^O_2_ pathways.

As the *fc2, flu*, and *ch1* mutants produce ^1^O_2_ within chloroplasts that leads to retrograde signaling (to control similar sets of genes), cell death, and (at least in *flu* and *fc2*) chloroplast degradation, we were initially surprised multiple pathways can be activated by this specific ROS. One possibility we suggest that the exact location of ^1^O_2_ production determines which signal is activated. In *flu* mutants, this likely occurs in the thylakoid grana margins where the EX proteins localize (Dogra et al. 2022; Wang et al. 2016). These grana margins are the site of PSII repair and tetrapyrrole synthesis, both potential sources of ^1^O_2_ in the light. On the other hand, *ch1* mutants produce ^1^O_2_ within the grana core, the site of active PSII (Ramel et al. 2013). Researchers have not yet determined the exact site of ^1^O_2_ production in *fc2* and *acd2* mutants. However, some work suggests Proto accumulates in the chloroplast envelope and stromal fractions of pea and beet, respectively (Mohapatra et al. 2002; Mohapatra et al. 2007). Thus, ^1^O_2_ in *fc2* may represent a more advanced stage of photo-oxidative stress where damage has spread throughout the chloroplast.

Another possibility for multiple pathways could be that the kinetics of ^1^O_2_ accumulation affects signaling. ^1^O_2_ is produced almost instantly after light exposure in *flu* mutants due to the accumulation of Pchlide in the dark (Meskauskiene et al. 2001). On the other hand, ^1^O_2_ production in the other mutants is slower and accumulates over time (Ramel et al. 2013; Woodson et al. 2015). The identification of additional signaling components should help resolve these possibilities.

Ultimately, we do not yet know why chloroplasts require multiple ^1^O_2_ signaling pathways to respond to stress or what the roles of these pathways are under natural environmental stresses. However, some experiments have offered clues. Under severe photo-inhibitory conditions that lead to bleaching and cell death, the *fc2*/*ch1* pathway may prevail. *pub4-6* (shown here) and *oxi1* (shown here and (Shumbe et al. 2016)) mitigate cell death under EL stress. We have demonstrated other suppressors of *fc2* cell death slow light-induced photo-bleaching in leaves (i.e., *ppr30, mterf9* (Alamdari et al. 2020) and cotyledons (i.e. *ctps2*) (Alamdari et al. 2021)). Under milder, non-photo-inhibitory light stress, plants may utilize the *flu* pathway. *ex1* blocks the formation of microlesions under moderate light stress that did not associate with non-enzymatic lipid peroxidation (Kim et al. 2012). Furthermore, these pathways could integrate different types of stress signals. Prior work links PUB4 and OXI1 to basal defense pathways (Rentel et al. 2004; Wang et al. 2022), and PUB4 plays a role in nitrogen and carbon starvation (Kikuchi et al. 2020). These findings suggest the *fc2*/*ch1* pathway might integrate with defense and senescence pathways. On the other hand, researchers have shown EX1 has a role in systemic acquired acclimation responses to EL stress (Carmody et al. 2016), suggesting plants use the *flu* pathway for distal signaling. These results along with our current findings demonstrate that chloroplast ^1^O_2_ signaling is a complex process requiring additional investigation. However, the existence of two (or more) chloroplast pathways may allow plants to better respond to their surroundings and to thrive in stressful and dynamic environments.

## Funding

The authors acknowledge the Division of Chemical Sciences, Geosciences, and Biosciences, Office of Basic Energy Sciences of the U.S. Department of Energy grant DE-SC0019573 awarded to JDW. DWT was supported by the University of Arizona University Fellows Award, the John Boynton Fellowship (School of Plant Sciences), and the Graduate College Completion Fellowship. MAK was supported by the University of Arizona University Fellows Award.

## Authors’ contributions

DWT, MAK, and JDW planned and designed the research. DWT performed all genetic analyses, metabolite measurements, meta-data analyses, and physiological growth experiments. DWT and RAE performed all genotyping and gene expression experiments. MAK performed all EL growth experiments and photosynthetic measurements. DWT and JDW performed the SOSG assays. JDW conceived the original scope of the project and managed the project. DWT and JDW wrote the manuscript. All authors contributed to data analysis, collection, and interpretation.

Acknowledgments

The authors wish to thank Matthew Lemke (University of Arizona) for technical assistance with trypan blue staining, Emmanuel Gonzalez (University of Arizona) for technical assistance with genotyping, and Kamran Alamdari (University of Arizona) for technical assistance propagating mutant lines.

## Short legends for supporting information

### Supplemental Figures

Fig. S1. Meta-analysis of transcriptome expression data from three chloroplast ^1^O_2_-producing mutant backgrounds. Fig. S2. GO Term analysis for ^1^O_2_-producing genetic backgrounds.

Fig. S3. GO Term analysis for transcriptome overlap between ^1^O_2_-producing genetic backgrounds and ß-cyclocitral treatment.

Fig. S4. GO Term analysis for transcriptome overlap between ^1^O_2_-producing genetic backgrounds and ESORGs. Fig. S5. Effects of *ex1* and *ex2* mutations on cell death in the *fc2* mutant background.

Fig. S6. *EX2* expression levels in *ex2* T-DNA mutants.

Fig. S7. Analysis of extended dark periods on retrograde signaling and the severity of cell death in *flu* mutant seedlings. Fig. S8. Effect of singlet oxygen signaling mutations on EL-induced phenotypes.

Fig. S9. The *pub4-6* mutation slows the progression of spontaneous cell death in the *acd2* mutant

## Supplementary Tables

Table S1. Mutant lines used in study

Table S2. Primes used for RT-qPCR and genotyping

Table S3. Nuclear-encoded genes included on the Affymetrix GeneChip Arabidopsis ATH1 Genome Array Tables S4. Genes included on the CATv5 microarray (Complete Arabidopsis Transcriptome Microarray)

Table S5. Nuclear-encoded genes shared by the Affymetrix GeneChip Arabidopsis ATH1 and CATv5 Microarrays Table S6. Differentially expressed genes in the *fc2* mutant dataset

Table S7. Differentially expressed genes in the *flu* mutant dataset Table S8. Differentially expressed genes in the *ch1* mutant dataset

Table S9. Differentially expressed genes in the ß-cyclocitral treatment dataset Table S10. Singlet Oxygen Early Response Genes (ESORGS)

Table S11. Up-regulated genes shared by the three singlet oxygen producing mutants Table S12. Down-regulated genes shared by the three singlet oxygen producing mutants Table S13. Up-regulated genes shared between mutants and ß -cyclocitral treatment Table S14. Down-regulated genes shared between mutants and ß -cyclocitral treatment Table S15. Up-regulated ESORGS shared between mutant backgrounds

Table S16. Conditions for previously published transcript profiling experiments

Table S17. Common genes differentially expressed between *fc2, flu*, and *ch1* datasets. Table S18. Common genes differentially expressed by treatment with ß -cyclocitral Table S19. *Early Singlet Oxygen Response Genes* (*ESORGs*) induced mutants

Table S20. Gene ontology analysis of unique up-regulated genes from mutants

## Statements and Declarations

The authors report no conflict of interest

## Notes

### Competing Interest Statement

The authors have declared no competing interest.

## References

1. Alamdari K, Fisher KE, Sinson AB, Chory J, Woodson JD (2020) Roles for the chloroplast-localized PPR Protein 30 and the “Mitochondrial” Transcription Termination Factor 9 in chloroplast quality control. Plant J 103:735–751

2. Alamdari K, Fisher KE, Tano DW, Rai S, Palos KR, Nelson ADL, Woodson JD (2021) Chloroplast quality control pathways are dependent on plastid DNA synthesis and nucleotides provided by cytidine triphosphate synthase two. New Phytol 231:1431–1448

3. Albrecht V, Ingenfeld A, Apel K (2008) Snowy cotyledon 2: the identification of a zinc finger domain protein essential for chloroplast development in cotyledons but not in true leaves. Plant Mol Biol 66:599–608

4. Alonso JM, Stepanova AN, Leisse TJ, Kim CJ, Chen H, Shinn P, Stevenson DK, Zimmerman J, Barajas P, Cheuk R, Gadrinab C, Heller C, Jeske A, Koesema E, Meyers CC, Parker H, Prednis L, Ansari Y, Choy N, Deen H, Geralt M, Hazari N, Hom E, Karnes M, Mulholland C, Ndubaku R, Schmidt I, Guzman P, Aguilar-Henonin L, Schmid M, Weigel D, Carter DE, Marchand T, Risseeuw E, Brogden D, Zeko A, Crosby WL, Berry CC, Ecker JR (2003) Genome-wide insertional mutagenesis of Arabidopsis thaliana. Science 301:653–7

5. Apel K, Hirt H (2004) Reactive oxygen species: metabolism, oxidative stress, and signal transduction. Annu Rev Plant Biol 55:373–399

6. Asada K (2006) Production and scavenging of reactive oxygen species in chloroplasts and their functions. Plant physiology 141:391–396

7. Barkan A, Small I (2014) Pentatricopeptide repeat proteins in plants. Annu Rev Plant Biol 65:415–42

8. Baruah A, Simkova K, Apel K, Laloi C (2009) Arabidopsis mutants reveal multiple singlet oxygen signaling pathways involved in stress response and development. Plant Mol Biol 70:547–63

9. Boyle EI, Weng S, Gollub J, Jin H, Botstein D, Cherry JM, Sherlock G (2004) GO::TermFinder—open source software for accessing Gene Ontology information and finding significantly enriched Gene Ontology terms associated with a list of genes. Bioinformatics 20:3710–3715

10. Bruggemann E, Handwerger K, Essex C, Storz G (1996) Analysis of fast neutron-generated mutants at the Arabidopsis thaliana HY4 locus. Plant J 10:755–60

11. Callis J (2014) The ubiquitination machinery of the ubiquitin system. The Arabidopsis book / American Society of Plant Biologists 12:e0174

12. Camehl I, Drzewiecki C, Vadassery J, Shahollari B, Sherameti I, Forzani C, Munnik T, Hirt H, Oelmüller R (2011) The OXI1 kinase pathway mediates Piriformospora indica-induced growth promotion in Arabidopsis. PLoS Pathog 7:e1002051

13. Carmody M, Crisp PA, d’Alessandro S, Ganguly D, Gordon M, Havaux M, Albrecht-Borth V, Pogson BJ (2016) Uncoupling High Light Responses from Singlet Oxygen Retrograde Signaling and Spatial-Temporal Systemic Acquired Acclimation. Plant Physiol 171:1734–49

14. Chan KX, Mabbitt PD, Phua SY, Mueller JW, Nisar N, Gigolashvili T, Stroeher E, Grassl J, Arlt W, Estavillo GM, Jackson CJ, Pogson BJ (2016) Sensing and signaling of oxidative stress in chloroplasts by inactivation of the SAL1 phosphoadenosine phosphatase. Proc Natl Acad Sci U S A 113:E4567–76

15. Chan KX, Phua SY, Crisp P, McQuinn R, Pogson BJ (2015) Learning the Languages of the Chloroplast: Retrograde Signaling and Beyond. Annu Rev Plant Biol 67:25–53

16. Chen S, Kim C, Lee JM, Lee HA, Fei Z, Wang L, Apel K (2015) Blocking the QB-binding site of photosystem II by tenuazonic acid, a non-host-specific toxin of Alternaria alternata, activates singlet oxygen-mediated and EXECUTER-dependent signalling in Arabidopsis. Plant Cell Environ 38:1069–80

17. Choudhury FK, Rivero RM, Blumwald E, Mittler R (2017) Reactive oxygen species, abiotic stress and stress combination. Plant J 90:856–867

18. D’Alessandro S, Beaugelin I, Havaux M (2020) Tanned or Sunburned: How Excessive Light Triggers Plant Cell Death. Molecular Plant 13:1545–1555

19. Danon A, Coll NS, Apel K (2006) Cryptochrome-1-dependent execution of programmed cell death induced by singlet oxygen in Arabidopsis thaliana (vol 103, pg 17036, 2006). Proceedings of the National Academy of Sciences of the United States of America 103:18875–18875

20. Dogra V, Duan J, Lee KP, Lv S, Liu R, Kim C (2017) FtsH2-Dependent Proteolysis of EXECUTER1 Is Essential in Mediating Singlet Oxygen-Triggered Retrograde Signaling in Arabidopsis thaliana. Front Plant Sci 8:1145

21. Dogra V, Kim C (2019) Singlet Oxygen Metabolism: From Genesis to Signaling. Front Plant Sci 10:1640

22. Dogra V, Li M, Singh S, Li M, Kim C (2019) Oxidative post-translational modification of EXECUTER1 is required for singlet oxygen sensing in plastids. Nat Commun 10:2834

23. Dogra V, Singh RM, Li M, Li M, Singh S, Kim C (2022) EXECUTER2 modulates the EXECUTER1 signalosome through its singlet oxygen-dependent oxidation. Mol Plant 15:438–453

24. Fisher KE, Krishnamoorthy P, Joens MS, Chory J, Fitzpatrick JAJ, Woodson JD (2022) Singlet Oxygen Leads to Structural Changes to Chloroplasts During their Degradation in the Arabidopsis thaliana plastid ferrochelatase two Mutant. Plant and Cell Physiology 63:248–264

25. Foyer CH (2018) Reactive oxygen species, oxidative signaling and the regulation of photosynthesis. Environ Exp Bot 154:134–142

26. Havaux M, Dall’osto L, Bassi R (2007) Zeaxanthin has enhanced antioxidant capacity with respect to all other xanthophylls in Arabidopsis leaves and functions independent of binding to PSII antennae. Plant Physiol 145:1506–20

27. Jeran N, Rotasperti L, Frabetti G, Calabritto A, Pesaresi P, Tadini L (2021) The PUB4 E3 Ubiquitin Ligase Is Responsible for the Variegated Phenotype Observed upon Alteration of Chloroplast Protein Homeostasis in Arabidopsis Cotyledons. Genes (Basel) 12:

28. Kikuchi Y, Nakamura S, Woodson JD, Ishida H, Ling Q, Hidema J, Jarvis RP, Hagihara S, Izumi M (2020) Chloroplast Autophagy and Ubiquitination Combine to Manage Oxidative Damage and Starvation Responses. Plant Physiol 183:1531–1544

29. Kim C, Meskauskiene R, Zhang S, Lee KP, Lakshmanan Ashok M, Blajecka K, Herrfurth C, Feussner I, Apel K (2012) Chloroplasts of Arabidopsis are the source and a primary target of a plant-specific programmed cell death signaling pathway. The Plant cell 24:3026–39

30. Kleinboelting N, Huep G, Kloetgen A, Viehoever P, Weisshaar B (2012) GABI-Kat SimpleSearch: new features of the Arabidopsis thaliana T-DNA mutant database. Nucleic acids research 40:D1211–5

31. Lee KP, Kim C, Landgraf F, Apel K (2007) EXECUTER1- and EXECUTER2-dependent transfer of stress-related signals from the plastid to the nucleus of Arabidopsis thaliana. Proc Natl Acad Sci U S A 104:10270–5

32. Lemke MD, Fisher EM, Kozlowska MA, Tano DW, Woodson JD (2021) The core autophagy machinery is not required for chloroplast singlet oxygen-mediated cell death in the Arabidopsis plastid ferrochelatase two mutant. BMC Plant Biology 21:342

33. Lu Y, Yao J (2018) Chloroplasts at the Crossroad of Photosynthesis, Pathogen Infection and Plant Defense. Int J Mol Sci 19:3900

34. Mach JM, Castillo AR, Hoogstraten R, Greenberg JT (2001) The Arabidopsis-accelerated cell death gene ACD2 encodes red chlorophyll catabolite reductase and suppresses the spread of disease symptoms. Proc Natl Acad Sci U S A 98:771–6

35. Meskauskiene R, Nater M, Goslings D, Kessler F, op den Camp R, Apel K (2001) FLU: a negative regulator of chlorophyll biosynthesis in Arabidopsis thaliana. Proc Natl Acad Sci U S A 98:12826–31

36. Mittler R (2017) ROS Are Good. Trends Plant Sci 22:11–19

37. Mohapatra A, Tripathy BC (2002) Detection of protoporphyrin IX in envelope membranes of pea chloroplasts. Biochem Biophys Res Commun 299:751–4

38. Mohapatra A, Tripathy BC (2007) Differential distribution of chlorophyll biosynthetic intermediates in stroma, envelope and thylakoid membranes in Beta vulgaris. Photosynth Res 94:401–10

39. Noctor G, Mhamdi A, Foyer CH (2014) The roles of reactive oxygen metabolism in drought: not so cut and dried. Plant Physiol 164:1636–48

40. Noctor G, Reichheld JP, Foyer CH (2018) ROS-related redox regulation and signaling in plants. Semin Cell Dev Biol 80:3–12

41. Ogilby PR (2010) Singlet oxygen: there is indeed something new under the sun. Chemical Society reviews 39:3181–209

42. Oliveros JC ((2007-2015)) Venny. An interactive tool for comparing lists with Venn’s diagrams. https://bioinfogp.cnb.csic.es/tools/venny/index.html

43. op den Camp RG, Przybyla D, Ochsenbein C, Laloi C, Kim C, Danon A, Wagner D, Hideg E, Gobel C, Feussner I, Nater M, Apel K (2003) Rapid induction of distinct stress responses after the release of singlet oxygen in Arabidopsis. Plant Cell 15:2320–2332

44. Page MT, Kacprzak SM, Mochizuki N, Okamoto H, Smith AG, Terry MJ (2017) Seedlings Lacking the PTM Protein Do Not Show a genomes uncoupled (gun) Mutant Phenotype. Plant Physiol 174:21–26

45. Page MT, McCormac AC, Smith AG, Terry MJ (2017) Singlet oxygen initiates a plastid signal controlling photosynthetic gene expression. New Phytol 213:1168–1180

46. Papenbrock J, Mishra S, Mock HP, Kruse E, Schmidt EK, Petersmann A, Braun HP, Grimm B (2001) Impaired expression of the plastidic ferrochelatase by antisense RNA synthesis leads to a necrotic phenotype of transformed tobacco plants. Plant J 28:41–50

47. Pattanayak GK, Venkataramani S, Hortensteiner S, Kunz L, Christ B, Moulin M, Smith AG, Okamoto Y, Tamiaki H, Sugishima M, Greenberg JT (2012) Accelerated cell death 2 suppresses mitochondrial oxidative bursts and modulates cell death in Arabidopsis. Plant J 69:589–600

48. Pospíšil P (2016) Production of Reactive Oxygen Species by Photosystem II as a Response to Light and Temperature Stress. Frontiers in Plant Science 7:

49. Pruzinská A, Anders I, Aubry S, Schenk N, Tapernoux-Lüthi E, Müller T, Kräutler B, Hörtensteiner S (2007) In vivo participation of red chlorophyll catabolite reductase in chlorophyll breakdown. Plant Cell 19:369–87

50. Ramel F, Birtic S, Ginies C, Soubigou-Taconnat L, Triantaphylides C, Havaux M (2012) Carotenoid oxidation products are stress signals that mediate gene responses to singlet oxygen in plants. Proceedings of the National Academy of Sciences of the United States of America 109:5535–40

51. Ramel F, Ksas B, Akkari E, Mialoundama AS, Monnet F, Krieger-Liszkay A, Ravanat JL, Mueller MJ, Bouvier F, Havaux M (2013) Light-induced acclimation of the Arabidopsis chlorina1 mutant to singlet oxygen. The Plant cell 25:1445–62

52. Rentel MC, Lecourieux D, Ouaked F, Usher SL, Petersen L, Okamoto H, Knight H, Peck SC, Grierson CS, Hirt H, Knight MR (2004) OXI1 kinase is necessary for oxidative burst-mediated signalling in Arabidopsis. Nature 427:858–61

53. Rosenwasser S, Fluhr R, Joshi JR, Leviatan N, Sela N, Hetzroni A, Friedman H (2013) ROSMETER: a bioinformatic tool for the identification of transcriptomic imprints related to reactive oxygen species type and origin provides new insights into stress responses. Plant Physiol 163:1071–83

54. Shin J, Kim K, Kang H, Zulfugarov IS, Bae G, Lee CH, Lee D, Choi G (2009) Phytochromes promote seedling light responses by inhibiting four negatively-acting phytochrome-interacting factors. Proceedings of the National Academy of Sciences of the United States of America 106:7660–5

55. Shumbe L, Bott R, Havaux M (2014) Dihydroactinidiolide, a high light-induced beta-carotene derivative that can regulate gene expression and photoacclimation in Arabidopsis. Mol Plant 7:1248–51

56. Shumbe L, Chevalier A, Legeret B, Taconnat L, Monnet F, Havaux M (2016) Singlet Oxygen-Induced Cell Death in Arabidopsis under High-Light Stress Is Controlled by OXI1 Kinase. Plant Physiol 170:1757–71

57. Shumbe L, D’Alessandro S, Shao N, Chevalier A, Ksas B, Bock R, Havaux M (2017) METHYLENE BLUE SENSITIVITY 1 (MBS1) is required for acclimation of Arabidopsis to singlet oxygen and acts downstream of beta-cyclocitral. Plant Cell Environ 40:216–226

58. Suo J, Zhao Q, David L, Chen S, Dai S (2017) Salinity Response in Chloroplasts: Insights from Gene Characterization. Int J Mol Sci 18:1011

59. Supek F, Bošnjak M, Škunca N, Šmuc T (2011) REVIGO summarizes and visualizes long lists of gene ontology terms. PLoS One 6:e21800

60. Triantaphylides C, Krischke M, Hoeberichts FA, Ksas B, Gresser G, Havaux M, Van Breusegem F, Mueller MJ (2008) Singlet oxygen is the major reactive oxygen species involved in photooxidative damage to plants. Plant physiology 148:960–8

61. Uberegui E, Hall M, Lorenzo O, Schroder WP, Balsera M (2015) An Arabidopsis soluble chloroplast proteomic analysis reveals the participation of the Executer pathway in response to increased light conditions. J Exp Bot 66:2067–77

62. Wagner D, Przybyla D, Op den Camp R, Kim C, Landgraf F, Lee KP, Wursch M, Laloi C, Nater M, Hideg E, Apel K (2004) The genetic basis of singlet oxygen-induced stress responses of Arabidopsis thaliana. Science 306:1183–5

63. Wang L, Kim C, Xu X, Piskurewicz U, Dogra V, Singh S, Mahler H, Apel K (2016) Singlet oxygen- and EXECUTER1-mediated signaling is initiated in grana margins and depends on the protease FtsH2. Proc Natl Acad Sci U S A 113:E3792–800

64. Wang LS, Leister D, Guan L, Zheng Y, Schneider K, Lehmann M, Apel K, Kleine T (2020) The Arabidopsis SAFEGUARD1 suppresses singlet oxygen-induced stress responses by protecting grana margins. Proceedings of the National Academy of Sciences of the United States of America 117:6918–6927

65. Wang Y, Wu Y, Zhong H, Chen S, Wong KB, Xia Y (2022) Arabidopsis PUB2 and PUB4 connect signaling components of pattern-triggered immunity. New Phytol 233:2249–2265

66. Warren CR (2008) Rapid Measurement of Chlorophylls with a Microplate Reader. Journal of Plant Nutrition 31:1231–1332

67. Wobbe L (2020) The molecular function of plant mTERFs as key regulators of organellar gene expression. Plant and Cell Physiology 61:2004–2017

68. Woodson JD (2019) Chloroplast stress signals: regulation of cellular degradation and chloroplast turnover. Curr Opin Plant Biol 52:30–37

69. Woodson JD (2022) Control of chloroplast degradation and cell death in response to stress. Trends Biochem Sci

70. Woodson JD, Joens MS, Sinson AB, Gilkerson J, Salome PA, Weigel D, Fitzpatrick JA, Chory J (2015) Ubiquitin facilitates a quality-control pathway that removes damaged chloroplasts. Science 350:450–4

71. Woodson JD, Perez-Ruiz JM, Chory J (2011) Heme synthesis by plastid ferrochelatase I regulates nuclear gene expression in plants. Curr Biol 21:897–903

