## Supplemental information for "Multiple pathways mediate chloroplast singlet oxygen stress signaling"

*Figure S1. Meta-analysis of transcriptome expression data from three chloroplast $^1O_2$-producing mutant backgrounds.*

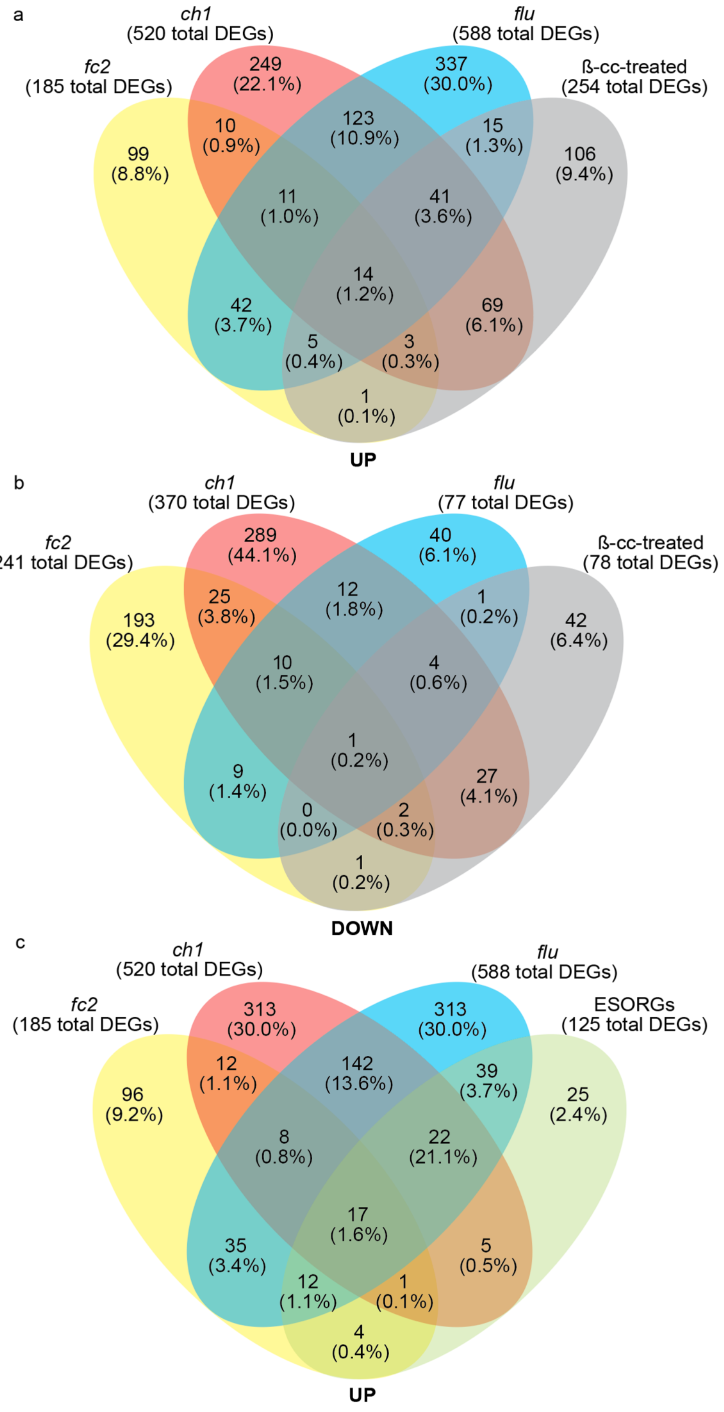


Venn diagram depicting common differentially expressed genes (DEGs) **A)** up-regulated or **B)** down-regulated during chloroplast $^1O_2$ stress in the *fc2* (Woodson et al. 2015), *flu* (op den Camp et al. 2003), and *ch1* (Ramel et al. 2013) genetic backgrounds compared to wt plants treated with β-cyclocitral (β-cc) (Ramel et al. 2012). **C)** Venn diagram showing common up-regulated genes during chloroplast $^{1}O_2$ stress in the *fc2* (Woodson et al. 2015), *flu* (op den Camp et al. 2003), and *ch1* (Ramel et al. 2013) genetic backgrounds compared to Early Singlet Oxygen Response Genes (ESORGs) in *flu* mutant seedlings (Dogra et al. 2017). The number of overlapping DEGs (and percent proportion to total DEGs in the analysis) is indicated within each colored area.

*Figure S2. GO Term analysis for $^1O_2$-producing genetic backgrounds.*

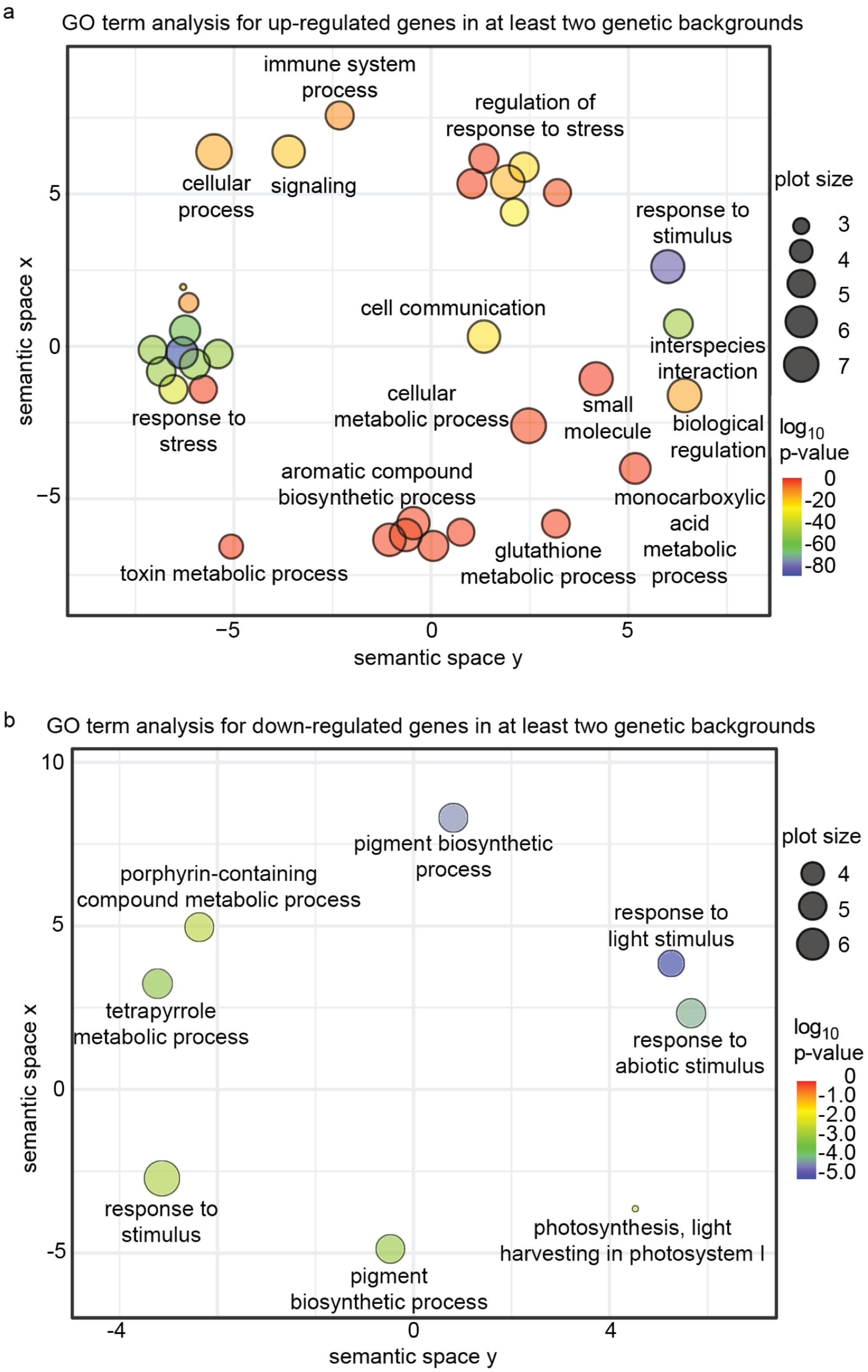


Scatterplot of gene ontology (GO) terms for differentially expressed genes shared among at least two chloroplast $^1O_2$ accumulating mutants; *fc2* (Woodson et al. 2015), *flu* (op den Camp et al.

2003), and *ch1* (Ramel et al. 2013). Shown are enriched GO terms from shared **A)** up-regulated or **B)** down-regulated genes. GO terms were identified using GO::TermFinder (https://go.princeton.edu/cgi-bin/GOTermFinder) (Boyle et al. 2004) and selected based on a p-value ≤ 0.01. Qualifying GO terms were exported to REVIGO ("small" option for filtering was applied) to create the scatterplot (http://revigo.irb.hr) (Supek et al. 2011), which shows the GO cluster representatives (i.e., remaining GO terms after removal of redundancies) as circles in a grid representing semantic similarities. The closer the circles are to each other, the more related the GO terms are. The size of the circle indicates the frequency of the GO term in the underlying GO annotation database. As such, larger circles represent more general terms. The color of the circles represents the $\log_{10}$ (p-value) of the GO term.

*Figure S3. GO Term analysis for transcriptome overlap between $^1O_2$-producing genetic backgrounds and β-cyclocitral treatment.*

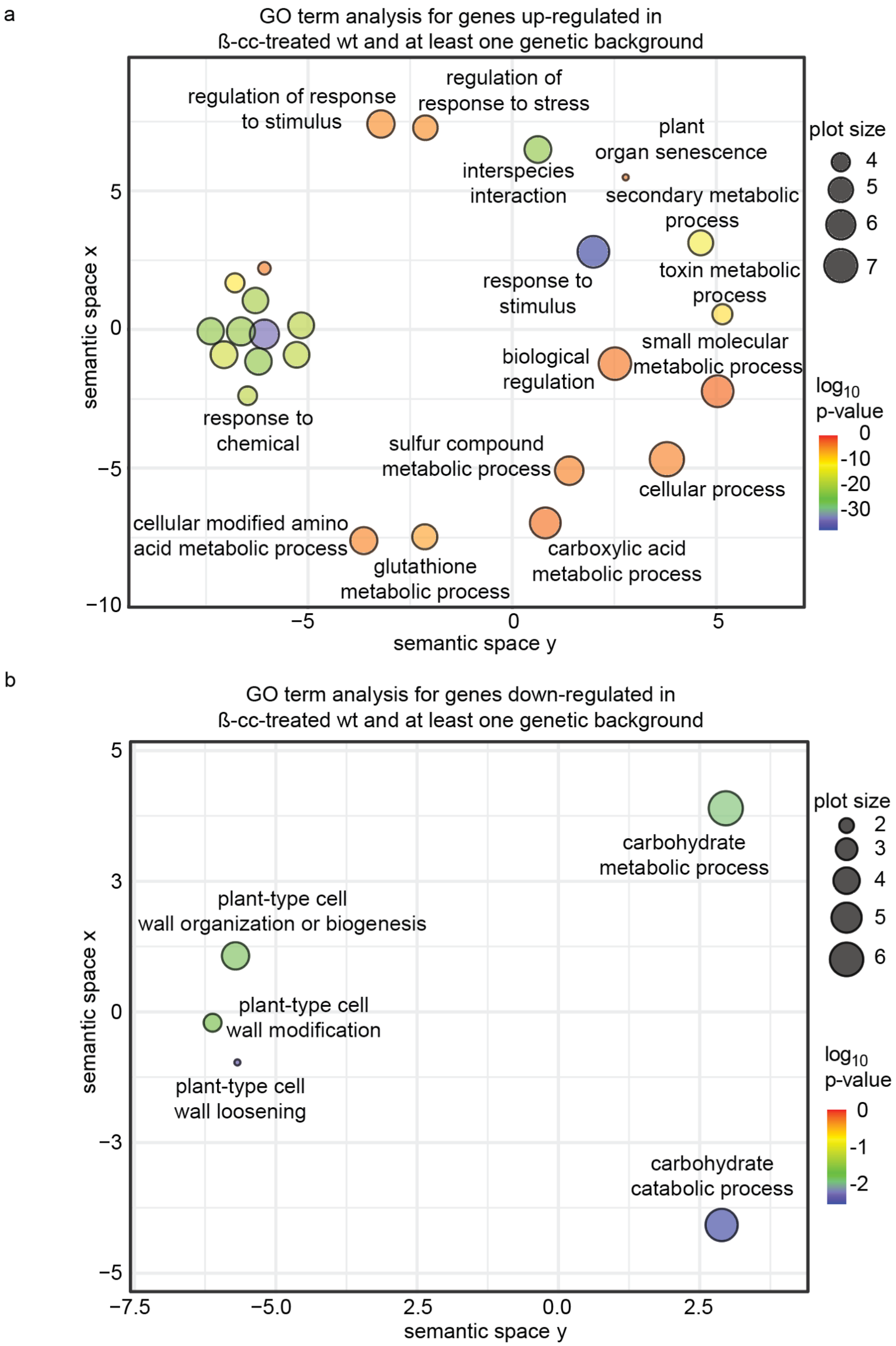


Scatterplot of gene ontology (GO) terms for differentially expressed genes shared among β-cyclocitral (β-cc)-treated wt plants (Ramel et al. 2012) and at least one chloroplast $^1O_2$

accumulating mutant; *fc2* (Woodson et al. 2015), *flu* (op den Camp et al. 2003), and *ch1* (Ramel et al. 2013). Shown are enriched GO terms from shared **A)** up-regulated or **B)** down-regulated genes. GO terms were identified using GO::TermFinder (https://go.princeton.edu/cgi-bin/GOTermFinder) (Boyle et al. 2004) and selected based on a p-value ≤ 0.01. Qualifying GO terms were exported to REVIGO ("small" option for filtering was applied) to create the scatterplot (http://revigo.irb.hr) (Supek et al. 2011), which shows the GO cluster representatives (i.e., remaining GO terms after removal of redundancies) as circles in a grid representing semantic similarities. The closer the circles are to each other, the more related the GO terms are. The size of the circle indicates the frequency of the GO term in the underlying GO annotation database. Aa such, larger circles represent more general terms. The color of the circles represents the $\log_{10}$ (p-value) of the GO term.

*Figure S4. GO Term analysis for transcriptome overlap between $^{1}O_2$-producing genetic backgrounds and ESORGS.*

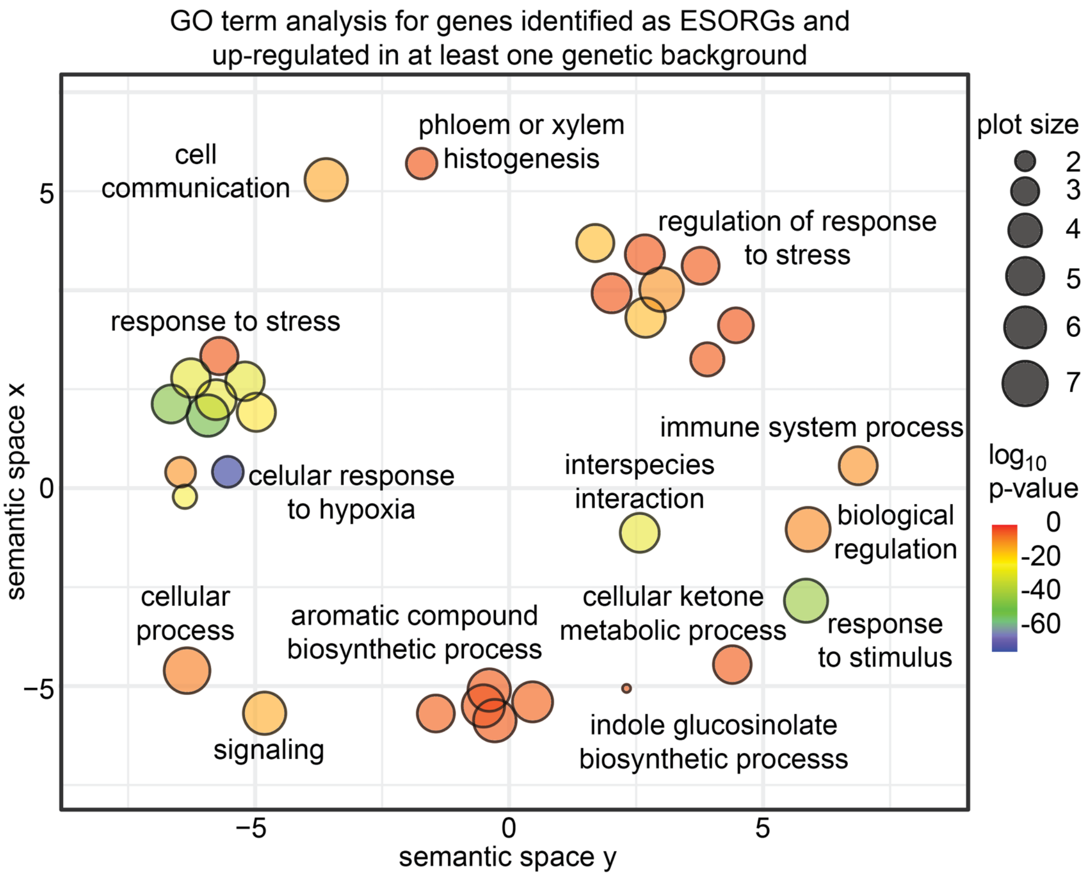


Scatterplot of gene ontology (GO) terms for Early Singlet Oxygen Response Genes (ESORGs) in *flu* seedlings (Dogra et al. 2017) found to be significantly induced in at least one chloroplast $^{1}O_2$ accumulating mutant; *fc2* (Woodson et al. 2015), *flu* (op den Camp et al. 2003), and *ch1* (Ramel et al. 2013). Shown are enriched GO terms from shared **A)** up-regulated or **B)** down-regulated genes. GO terms were identified using GO::TermFinder (https://go.princeton.edu/cgi-bin/GOTermFinder) (Boyle et al. 2004) and selected based on a p-value ≤ 0.01. Qualifying GO terms were exported to REVIGO ("small" option for filtering was applied) to create the scatterplot (http://revigo.irb.hr) (Supek et al. 2011), which shows the GO cluster representatives (i.e., remaining GO terms after removal of redundancies) as circles in a grid representing semantic similarities. The closer the circles are to each other, the more related the GO terms are. The size of the circle indicates the frequency of the GO term in the underlying GO annotation database. Aa such, larger circles represent more general terms. The color of the circles represents the $\log_{10}$ (p-value) of the GO term.

*Figure S5. Effects of ex1 and ex2 mutations on cell death in the fc2* mutant background.

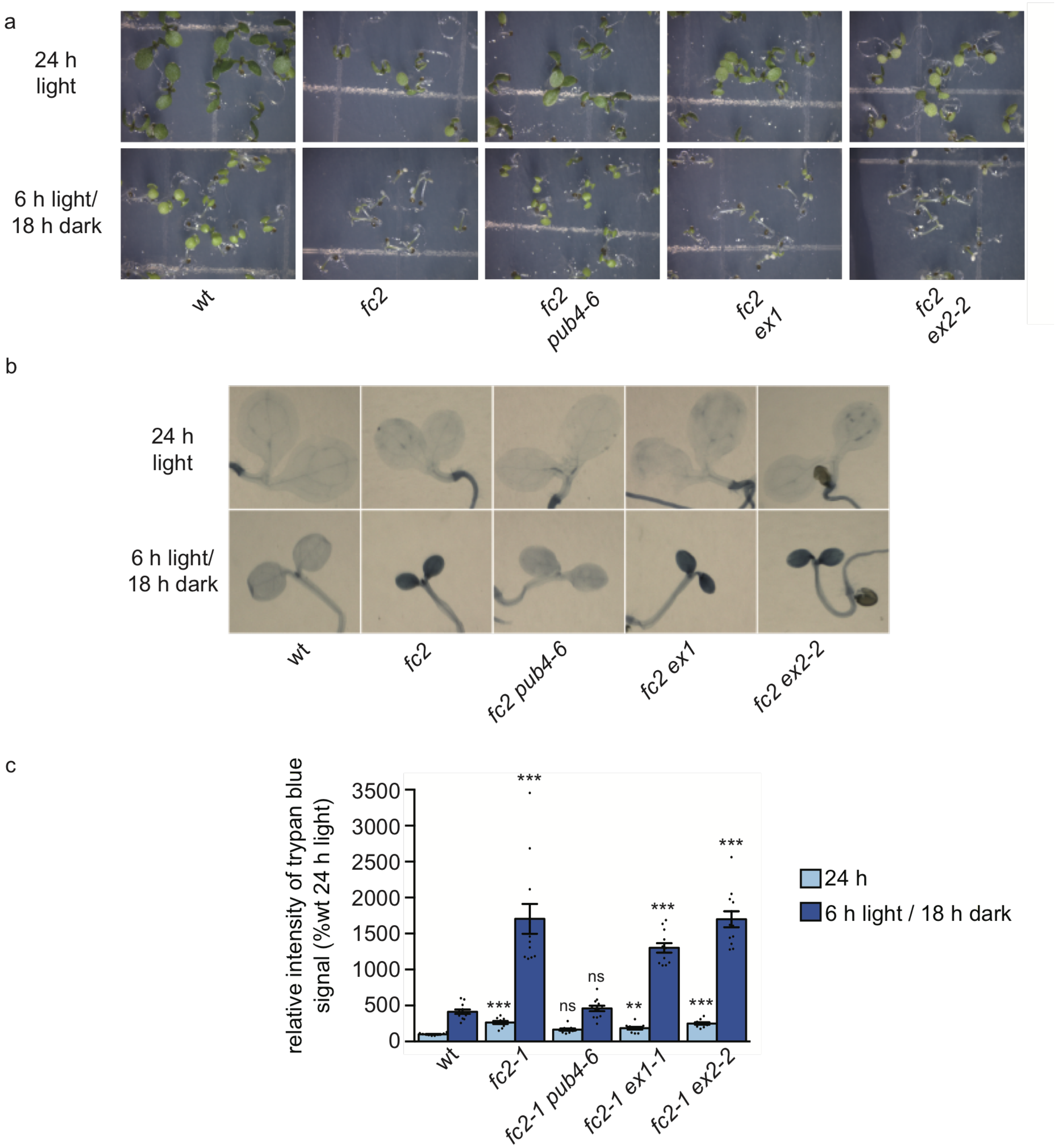


The effects of the *ex1* and *ex2* mutations on the *fc2* cell death phenotype were assessed. **A)** Shown are six-day-old seedlings grown under constant light (24h) or diurnal cycling light (6h light / 18h dark) conditions. **B)** Representative images of the same seedlings stained with trypan blue. The dark blue color indicates cell death. **C)** Shown are mean intensities of trypan blue signal (+/- SE, n ≥ 10 seedlings) from **B**. Statistical analyses were performed using a one-way ANOVA followed by a Tukey HSD test. Statistical significance in respect to *fc2* is indicated as follows: n.s. = p-value ≥ 0.05, *** = p-value ≤ 0.001. Closed circles represent individual data points.

*Figure S6. EX2 expression levels in ex2 T-DNA mutants.*

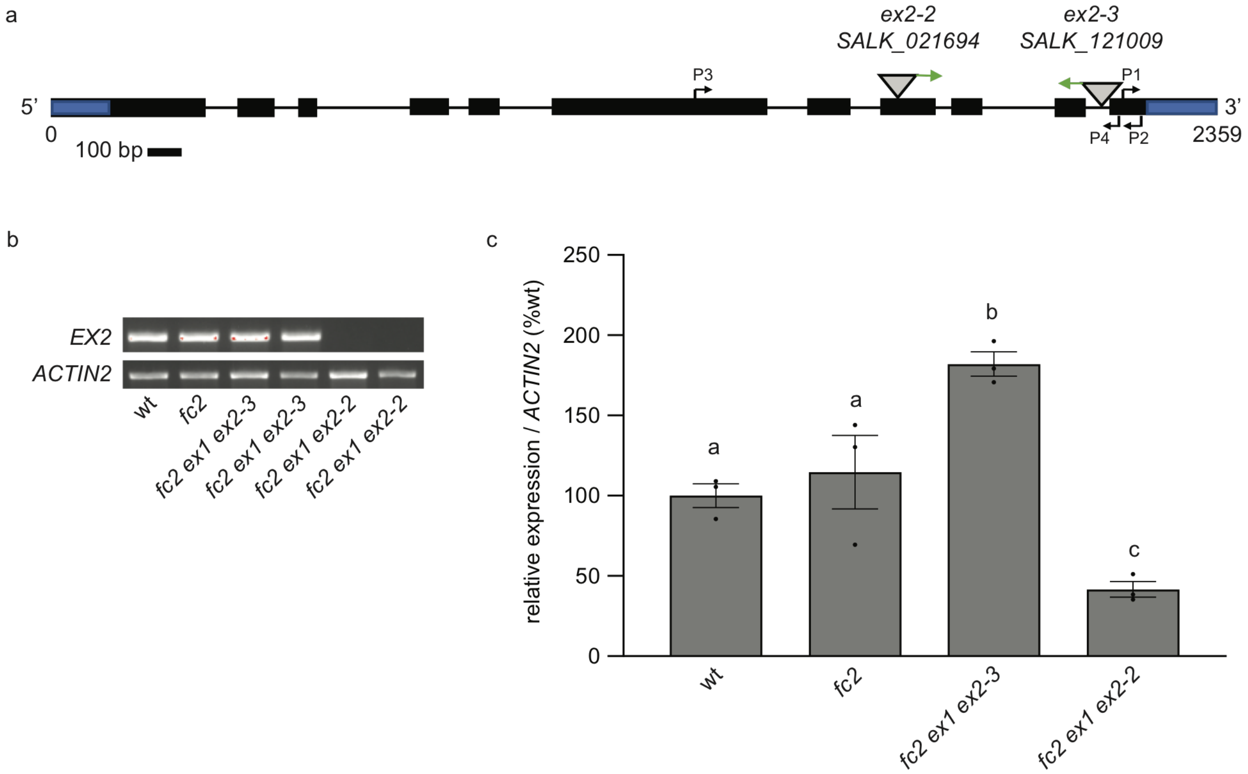


The effects of two *ex2* T-DNA mutant alleles were assessed. **A)** Gene diagram of *EX2* (*At1g27510*). Triangles represent T-DNA insertions with the right borders denoted by green arrows. The primers P1 and P2 are WLO1688/1689 used for RT-qPCR (panel **C** in this figure). The primers P3 and P4 are WLO1642/1643 used for semi-RT-qPCR (panel **B** in this figure, primer sequences in Table S2). The 5' and 3' untranslated regions are marked by a blue rectangle in the first and last exons respectively. Scale bar = 100bp. **B)** Shown is a semi-RT-qPCR analysis of cDNA generated from RNA extracted from five-day old seedlings grown in 24h constant light. PCR was repeated for 30 cycles. **C)** Shown is RT-qPCR analysis of cDNA generated from RNA extracted from four-day-old seedlings grown under 6h light / 18h dark cycling diurnal conditions (+/- SE, n = 3 seedling pools). Statistical analyses were performed using a one-way ANOVA followed by a Tukey HSD test. Different letters above bars indicate significant differences within data sets (p-value ≤ 0.05). Closed circles represent individual data points.

*Figure S7. Analysis of extended dark periods on retrograde signaling and the severity of cell death in flu mutant seedlings.*

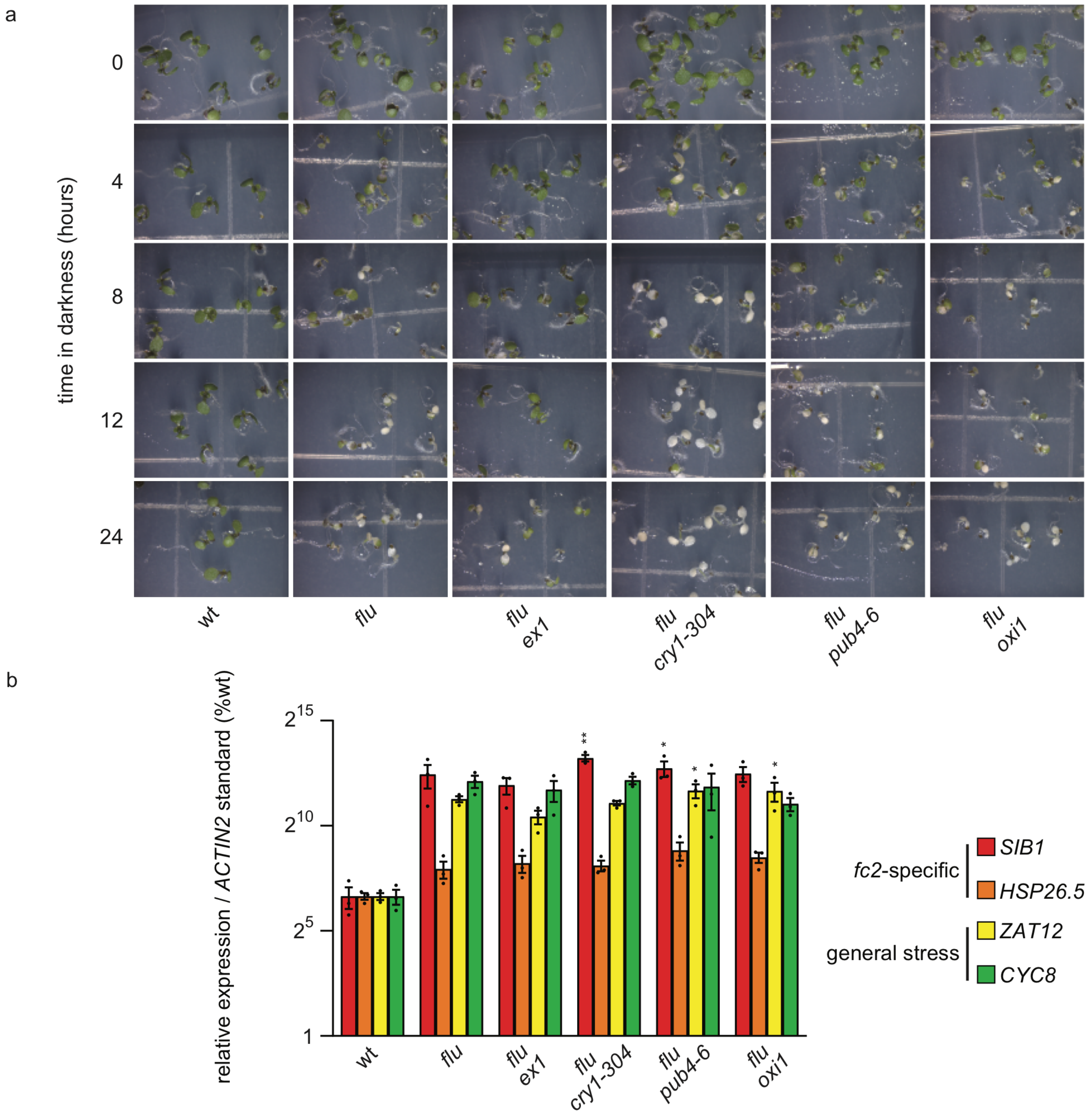


The extent of cotyledon bleaching in the *flu* mutant caused by increasing dark periods was tested. **A)** Shown are seedlings grown for five days in constant light, shifted to dark under the indicated amount of time (0-24 hours), and then shifted back to light for 36h (seven days total). Bleaching of cotyledons indicates cell death caused by $^1O_2$ signaling. The row for the 12h dark incubation was used in Fig. 4a. **B)** RT-qPCR of stress gene markers (from *fc2* mutants; *SIB1* and *HSP26.5* (Woodson et al. 2015)) and general stress (*ZAT12* and *CYC8* (Baruah et al. 2009)) of five-day old seedlings grown under 24h constant light then dark-incubated for 12 hours, harvested one hour after re-exposure to light. Shown are mean expression values (+/- SE, n = 3 biological replicates). Statistical analyses were performed using a one-way ANOVA followed by a Tukey HSD test. Statistical significance in respect to wt is indicated as follows: * = p-value ≤ 0.05, ** = p-value ≤ 0.01. Closed circles represent individual data points.

*Figure S8.* Effect of singlet oxygen signaling mutations on EL-induced phenotypes.

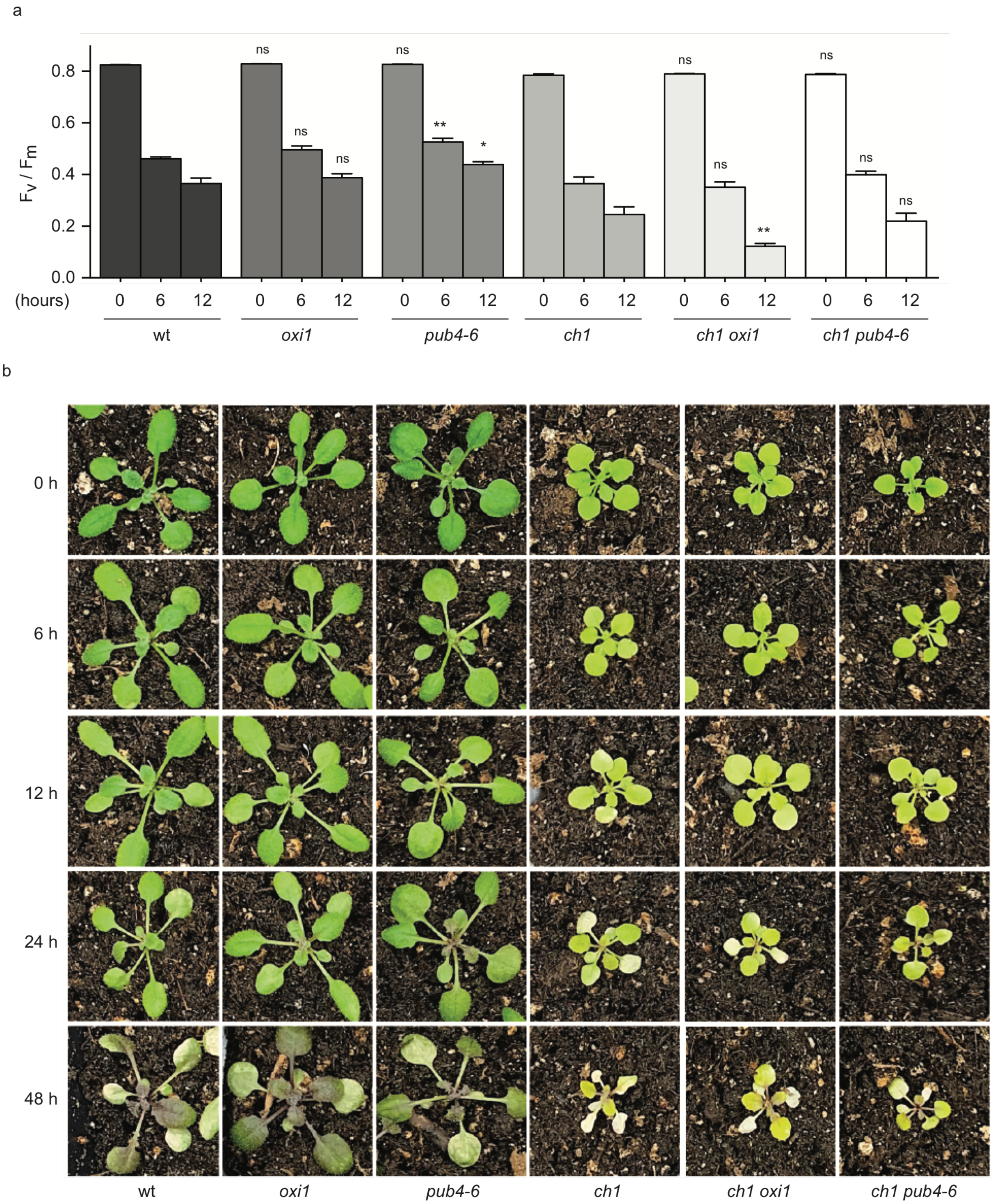


The effect of the *oxi1* and *pub4-6* mutations were tested in adult plants treated with excess light (EL). **A)** Time course analysis of maximum PSII quantum efficiency ($F_v/F_m$) in these plants during the initial 12 hours of EL (1,300 μmol $sec^{-1}$ $m^{-2}$) and 10 degrees C. **B)** Shown are representative images of EL-treated plants at the indicated times.

<u>*Figure S9*. The *pub4-6* mutation slows the progression of spontaneous cell death in the *acd2* mutant.</u>

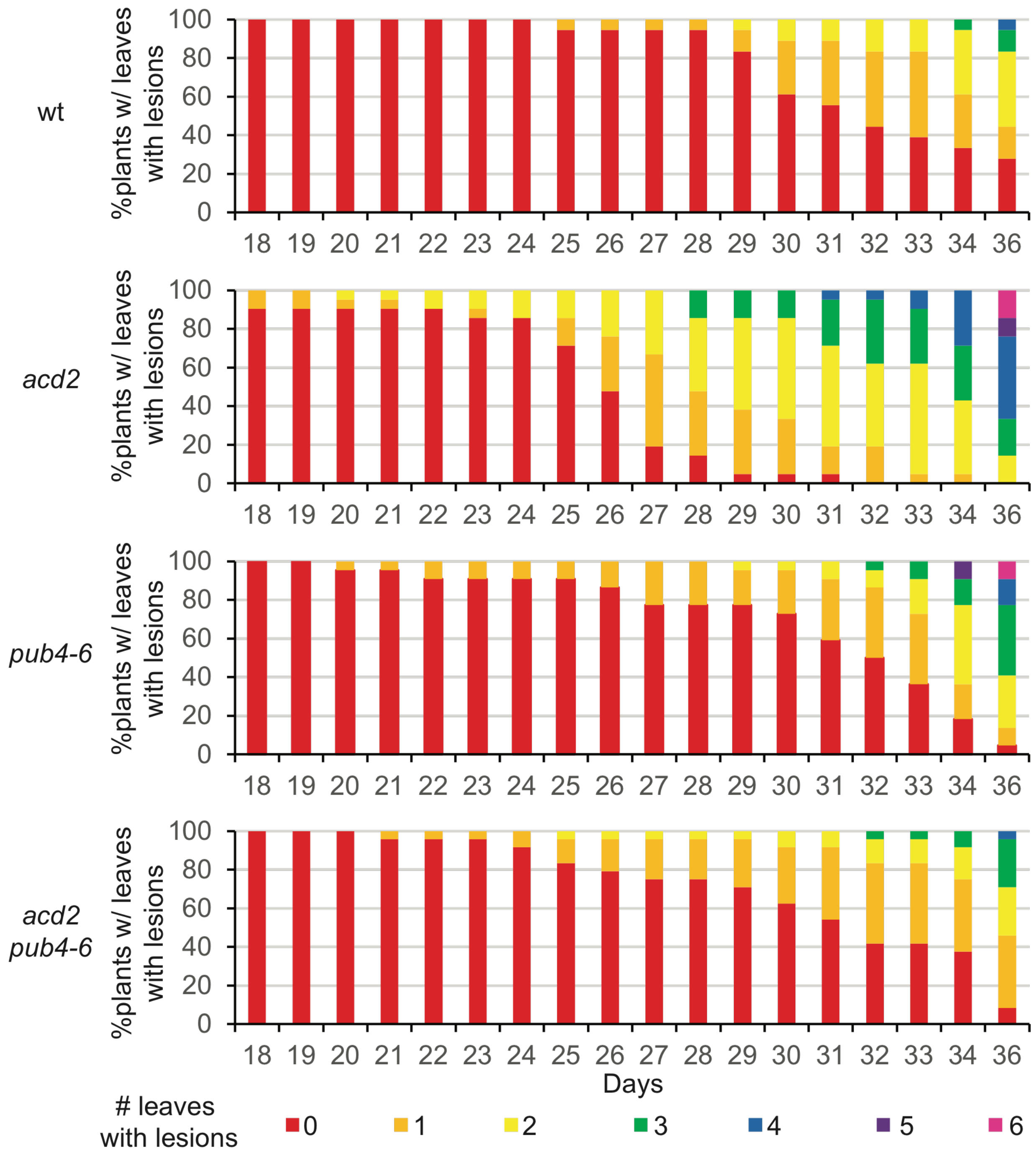


The *acd2* spontaneous cell death and lesion phenotypes were assessed. Shown are the percentage of plants containing different numbers of leaves with lesions from 18 to 36 days old (n ≥ 18 plants). Any type of discoloring was considered a lesion.

*Table S1. Mutant lines used in study*

| Mutant | Gene Name | Gene # | DNA/Protein Change | Notes | Ref. |
|---|---|---|---|---|---|
| *acd2-2* | *ACCELERATED CELL DEATH 2 ARABIDOPSIS THALIANA RED CHLOROPHYLL CATABOLITE REDUCTASE* | *At4g37000* | G to A substitution in the intronic splice site | Generated by ethyl methanesulfonate treatment | (Mach et al. 2001) |
| *ch1-1* | *CHLORINA 1*; *CHLOROPHYLL A OXYGENASE* | *At1g44446* | Deletion | Generated by X-ray Diffraction | (Havaux et al. 2007) |
| *cry1-304* | *CRYPTOCHROME 1* | *At4g08920* | Deletion | Generated by neutron mutagenesis | (Bruggemann et al. 1996) |
| *ex1* | *EXECUTER 1* | *At4g33630* | SALK_002088 T-DNA in 2$^{nd}$ exon | | (Lee et al. 2007) |
| *ex2-2* | *EXECUTER 2* | *At1g27510* | SALK_021694 T-DNA in 8$^{th}$ exon | | (Uberegui et al. 2015) |
| *ex2-3* | *EXECUTER 2* | *At1g27510* | SALK_121009 T-DNA in 10$^{th}$ intron | | This Study |
| *fc2-1* | *PLASTID FERROCHELATASE 2* | *At2g30390* | GABI_766H08<br>T-DNA in 5'UTR | Sulfadiazine Resistant | (Woodson et al. 2011) |
| *flu-1* | *FLUORESCENT IN BLUE LIGHT* | *At3g14110* | C-terminal amino acid substitution (A262V) | Generated by ethyl methanesulfonate treatment | (Meskauskiene et al. 2001) |
| *oxi1-1* | *OXIDATIVE SIGNAL-INDUCIBLE 1* | *At3g25250* | GABI_355H08 T-DNA in 2$^{nd}$ exon | | (Camehl et al. 2011) |
| *pub4-6* | *PLANT U-BOX 4* | *At2g23140* | U-box domain amino acid substitution (G255R) | Generated by ethyl methanesulfonate treatment | (Woodson et al. 2015) |

*Table S2. Primers used for RT-qPCR and genotyping*

| Gene | Primer orientation / name | Sequence |
|---|---|---|
| RT-qPCR primer pairs | | |
| *ACTIN2 / At3g18780* | For. / JP199 | GCACTTGCACCAAGCAGCAT |
| | Rev. / JP200 | CCTTTCAGGTGGTGCAACGAC |
| *SIB1 / At3g56710* | For. / JP589 | CAACCGGAGCCCATCTATT |
| | Rev. / JP590 | GGAGAAAGGTTGTGGTCGTC |
| *HSP26.5 / At1g52560* | For. / JP585 | CGAGCTTATCGTTGCCTGAT |
| | Rev. / JP586 | CTCCGCCTTAATGTCCTCAA |
| *BAP1 / At3g61190* | For. / JP338 | ATTGATGGATACGGTGGCCG |
| | Rev. / JP339 | CAGACCCCAAACCGGAACTC |
| *Atpase / At3g28580* | For. / JP336 | GAAGATCGGAAAAGCGTGGAA |
| | Rev. / JP337 | CCGGGTGGTCCAAACAAAAG |
| *ZAT12 / At5g59820* | For. / JP344 | GCGTTGGTTACACGCGCTT |
| | Rev. / JP345 | CTTCAACGTAGTCACCGTGGG |
| *CYC8 / At4g37370* | For. / JP1130 | AATGGGCATTGTCGAACGTG |
| | Rev. / JP1131 | TCGCCTTGTTCAATACATCCG |
| *NOD1 / At5g64870* | For. / JP340 | GCTGATGCTGCCTTCTATTCAA |
| | Rev. / JP341 | TGCGACAAGTCCCTCTGCA |
| *EX2 / At1g27510* | For. / WLO1688 | CATATGTGAAGGGCGCAGATC |
| | Rev. / WLO1689 | CAGACGGTTGAAAAGGACCAAG |
| Semi-qPCR primer pairs | | |
| *ACTIN2 / At3g18780* | For. /WLO1401 | GGCTGAGGCTGATGATATTC |
| | Rev. /WLO1402 | TCTGTGAACGATTCCTGGAC |
| *EX2 / At1g27510* | For. /WLO1642 | CGTGTGTCGGCAAATATAATGGA |
| | Rev. /WLO1643 | TCTGCGCCCTTCACATATGG |
| Genotyping primer pairs | | |

| | | |
|---|---|---|
| SALK T-DNA Left Border | LB1.3 | ATTTTGCCGATTTCGGAAC |
| GABI-KAT Left Border | Gabi-KAT 03144 | ATATTGACCATCATACTCATTGC |
| *acd2-2* dCAPS genotyping: *Pst*I digestion, wt = 124 bp, mutant = 149 bp | For. / JP1144 | GAGAATCTTAAAGTTTGTTTTGTTCTGCA |
| | Rev. / JP1145 | TCGACCACAAAGTTTGGAGCT |
| *cry1-304* | For. / WLO1451 | ATGTCTGGTTCTGTATCTGGTTGTGGTTC |
| | Rev. / WLO1452 | ATAGTTCTCATCCACAGCCCAAG |
| *ex1-1* (*SALK_002088*) | For. / JP415 | TCTGACACTTGATGGGAAAGG |
| | Rev. / JP416 | TAAGCTGCGACTCCTTTTCTG |
| *ex2-2* (*SALK_021694*) | For. / JP1140 | CACTAAGCTTGTCATCGGAGG |
| | Rev. / JP1141 | AAATGTCAATGTGGCTGGAAC |
| *ex2-3* (*SALK_121009*) | For. / JP1142 | TCCCTATTGATTTGCAGAAGC |
| | Rev. / JP1143 | ATTGTTTCCAGAGGAATTGGG |
| *fc2-1* (*GabiKat_766H08*) | For. / JP283 | GAGCAACGCCAAACATAGAAG |
| | Rev. / JP284 | TCAAAGGCAATGAATGTTTCC |
| *flu-1* dCAPS genotyping: *HpyI*88I digestion, wt = 127 bp, mutant = 102 bp | For. / JP1138 | AGCTTTGGAACTTGCCCAGA |
| | Rev. / JP1139 | TAGAACCATGGAGTGATACTGTCTG |
| *oxi1-1* (*Gabi_355H08*) | For. / JP1291 | CCTTTCCAAACAAAGCAAGTG |
| | Rev. / JP1292 | AAGAAACGTCTCTTCCGCTTC |
| *pub4-6* dCAPS genotyping: *Hpa*II digestion, wt = 97 bp, mutant = 121 bp | For. / JP742 | TATTAGAGTAGTGTGAGTCAGG |
| | Rev. / JP743 | GATCCAGTGATTGTGTCATCC |

*Table S16. Conditions for previously published transcript profiling experiments*

| Genotype | RNA Extraction | Expression Analysis | Data Sets Selected | Growing Conditions | Age | Ref. |
|---|---|---|---|---|---|---|
| *fc2* | RNeasy Plant Mini Kit (Qiagen) | Affymetrix GeneChip Arabidopsis ATH1 Genome Array. | 2 hr timepoint was used. *fc2* was compared to wt grown in the same conditions. | First grown in the dark for 4 days and then exposed to 2 hours of white light (120 µmol photons $m^{-2}$ $s^{-1}$). | 4-day old etiolated seedlings | (Woodson et al. 2015) |
| *flu* | Not listed | Affymetrix GeneChip Arabidopsis ATH1 Genome Array. | 1 hr timepoint was used. *flu* following dark/light shift was compared to wt treated similarly | First grown in continuous white light (80-100 photons µmol $m^{-2}$ $s^{-1}$). Incubated in dark for 8 hours and then exposed again to white light for 1 hr | Rosette stage adult plants | (op den Camp et al. 2003) |
| *ch1* | TRIzol extraction (Invitrogen) and Message Amp aRNA kit (Ambion) | CATMAv5<br>Complete Arabidopsis Transcriptome MicroArray. | Excess light treated *ch1* was compared to untreated *ch1*. | First grown in 8h white light (180 µmol photons $m^{-2}$ $s^{-1}$) / 16 h dark photoperiod. Then exposed to 2 days of 1,000 µmol photons $m^{-2}$ $s^{-1}$ (same photoperiod) | 5-8 week old adult plants | (Ramel et al. 2013) |

| | | | | | | |
|---|---|---|---|---|---|---|
| β-cc-treated wild type | TRIzol extraction (Invitrogen) and Message Amp aRNA kit (Ambion) | CATMAv5<br><br>Complete Arabidopsis Transcriptome MicroArray. | β-cc-treated wt was compared to untreated wt (mock treatment of distilled water). | First grown under a 16 hr white light (100 µmol photons $m^{-2}$ $s^{-1}$) / 8h dark photoperiod. Then exposed to 50 µL of β-CC for 4 hr in an airtight box (60 µmol $m^{-2}$ $s^{-1}$) | 4 week old adult plants | (Ramel et al. 2012) |
| *flu* RNAseq | RNeasy Plant Mini Kit (Qiagen) and NEBNext Ultra Directional RNA Library Prep Kit for Illumina | Libraries constructed using NEBNext Ultra Directional RNA Library Prep Kit for Illumina. Sequenced on an Illumina HiSeq 2500 platform to generate 100 bp paired-end reads. False discovery rate of 0.05 and 1 transcript/million minimum. | Light-exposed (both 30 and 60 minute time points) *flu* plants were compared to *flu* plants without reillumination. | First grown under continuous white light (40 µmol photons $m^{-2}$ $s^{-1}$) for 5 days and the incubated in the dark for 4 hrs. Next, seedlings were re-illuminated with white light for 30 and 60 min (both timepoints used in analysis) | 5-day old seedlings | (Dogra et al. 2017) |

*Table S17. Common genes differentially expressed between fc2, flu, and ch1 data sets*

| Genotype | Differentially expressed genes in data set (#) (total / unique) | Gene Overlap with *fc2* (# / %) | Gene Overlap with *flu* (# / %) | Gene Overlap with *ch1* (# / %) | Overlap among all 3 Genotypes (# / %) |
|---|---|---|---|---|---|
| Up-regulated Genes | | | | | |
| *fc2* | 185 / 100 | - | 72/38.9 | 38/20.5 | 25/13.5 |
| *flu* | 588 / 352 | 72/12.2 | - | 189/32.1 | 25/4.3 |
| *ch1* | 520 / 318 | 38/7.3 | 189/36.3 | - | 25/4.8 |
| Down-regulated Genes | | | | | |
| *fc2* | 241 / 194 | - | 20/8.3 | 38/15.8 | 11/4.6 |
| *flu* | 77 / 41 | 20/26.0 | - | 27/35.1 | 11/14.3 |
| *ch1* | 370 / 316 | 38/10.3 | 27/7.3 | - | 11/3.0 |
| Up- and Down-regulated Genes | | | | | |
| *fc2* | 426 / 294 | - | 92/21.6 | 76/17.8 | 36/8.5 |
| *flu* | 665 / 393 | 92/13.8 | - | 206/31.0 | 36/5.4 |
| *ch1* | 890 / 634 | 76/8.5 | 206/23.1 | - | 36/4.0 |

*Table S18. Common genes differentially expressed by treatment with β-cyclocitral.*

| Genotype | # of differentially expressed genes in common | # of differentially expressed genes overall | % Overlap |
|---|---|---|---|
| Up-regulated | | | |
| *fc2* | 23 | 185 | 12.4 |
| *flu* | 75 | 588 | 12.8 |
| *ch1* | 127 | 520 | 24.4 |
| Down-regulated | | | |
| *fc2* | 4 | 241 | 1.7 |
| *flu* | 6 | 77 | 7.8 |
| *ch1* | 34 | 370 | 9.2 |
| Up- and Down-regulated | | | |
| *fc2* | 34 | 426 | 8.0 |
| *flu* | 83 | 665 | 12.5 |
| *ch1* | 166 | 890 | 18.7 |

*Table S19. Early Singlet Oxygen Response Genes (ESORGs)* induced in mutants

| Genotype | # up-regulated ESORGs | # up-regulated genes | % Overlap |
|---|---|---|---|
| *fc2* | 34 | 185 | 8.0 |
| *flu* | 78 | 588 | 11.7 |
| *ch1* | 43 | 520 | 4.8 |

### References

1. Baruah A, Simkova K, Apel K, Laloi C (2009) Arabidopsis mutants reveal multiple singlet oxygen signaling pathways involved in stress response and development. Plant Mol Biol 70:547-63
2. Boyle EI, Weng S, Gollub J, Jin H, Botstein D, Cherry JM, Sherlock G (2004) GO::TermFinder—open source software for accessing Gene Ontology information and finding significantly enriched Gene Ontology terms associated with a list of genes. Bioinformatics 20:3710-3715
3. Bruggemann E, Handwerger K, Essex C, Storz G (1996) Analysis of fast neutron-generated mutants at the Arabidopsis thaliana HY4 locus. Plant J 10:755-60

4. Camehl I, Drzewiecki C, Vadassery J, Shahollari B, Sherameti I, Forzani C, Munnik T, Hirt H, Oelmüller R (2011) The OXI1 kinase pathway mediates Piriformospora indica-induced growth promotion in Arabidopsis. PLoS Pathog 7:e1002051
5. Dogra V, Duan J, Lee KP, Lv S, Liu R, Kim C (2017) FtsH2-Dependent Proteolysis of EXECUTER1 Is Essential in Mediating Singlet Oxygen-Triggered Retrograde Signaling in Arabidopsis thaliana. Front Plant Sci 8:1145
6. Havaux M, Dall'osto L, Bassi R (2007) Zeaxanthin has enhanced antioxidant capacity with respect to all other xanthophylls in Arabidopsis leaves and functions independent of binding to PSII antennae. Plant Physiol 145:1506-20
7. Lee KP, Kim C, Landgraf F, Apel K (2007) EXECUTER1- and EXECUTER2-dependent transfer of stress-related signals from the plastid to the nucleus of Arabidopsis thaliana. Proc Natl Acad Sci U S A 104:10270-5
8. Mach JM, Castillo AR, Hoogstraten R, Greenberg JT (2001) The Arabidopsis-accelerated cell death gene ACD2 encodes red chlorophyll catabolite reductase and suppresses the spread of disease symptoms. Proc Natl Acad Sci U S A 98:771-6
9. Meskauskiene R, Nater M, Goslings D, Kessler F, op den Camp R, Apel K (2001) FLU: a negative regulator of chlorophyll biosynthesis in Arabidopsis thaliana. Proc Natl Acad Sci U S A 98:12826-31
10. op den Camp RG, Przybyla D, Ochsenbein C, Laloi C, Kim C, Danon A, Wagner D, Hideg E, Gobel C, Feussner I, Nater M, Apel K (2003) Rapid induction of distinct stress responses after the release of singlet oxygen in Arabidopsis. Plant Cell 15:2320-2332
11. Ramel F, Birtic S, Ginies C, Soubigou-Taconnat L, Triantaphylides C, Havaux M (2012) Carotenoid oxidation products are stress signals that mediate gene responses to singlet oxygen in plants. Proceedings of the National Academy of Sciences of the United States of America 109:5535-40
12. Ramel F, Ksas B, Akkari E, Mialoundama AS, Monnet F, Krieger-Liszkay A, Ravanat JL, Mueller MJ, Bouvier F, Havaux M (2013) Light-induced acclimation of the Arabidopsis chlorina1 mutant to singlet oxygen. The Plant cell 25:1445-62
13. Supek F, Bošnjak M, Škunca N, Šmuc T (2011) REVIGO summarizes and visualizes long lists of gene ontology terms. PLoS One 6:e21800
14. Uberegui E, Hall M, Lorenzo O, Schroder WP, Balsera M (2015) An Arabidopsis soluble chloroplast proteomic analysis reveals the participation of the Executer pathway in response to increased light conditions. J Exp Bot 66:2067-77
15. Woodson JD, Joens MS, Sinson AB, Gilkerson J, Salome PA, Weigel D, Fitzpatrick JA, Chory J (2015) Ubiquitin facilitates a quality-control pathway that removes damaged chloroplasts. Science 350:450-4
16. Woodson JD, Perez-Ruiz JM, Chory J (2011) Heme synthesis by plastid ferrochelatase I regulates nuclear gene expression in plants. Curr Biol 21:897-903
